# EvANI benchmarking workflow for evolutionary distance estimation

**DOI:** 10.1101/2025.02.23.639716

**Authors:** Sina Majidian, Stephen Hwang, Mohsen Zakeri, Ben Langmead

## Abstract

Advances in long-read sequencing technology has led to a rapid increase in high-quality genome assemblies. These make it possible to compare genome sequences across the Tree of Life, deepening our understanding of evolutionary relationships. Average nucleotide identity (ANI) is a distance measure that has been applied to species delineation, building of guide trees, and searching large sequence databases. Since computing ANI is computationally expensive, the field has increasingly turned to sketch-based approaches that use assumptions and heuristics to speed this up. We propose a suite of simulated and real benchmark datasets, together with a rank-correlation-based metric, to study how these assumptions and heuristics impact distance estimates. We call this evaluation framework EvANI. With EvANI, we show that ANIb is the ANI estimation algorithm that best captures tree distance, though it is also the least efficient. We show that *k*-mer based approaches are extremely efficient and have consistently strong accuracy. We also show that some clades have inter-sequence distances that are best computed using multiple values of *k*, e.g. *k* = 10 and *k* = 19 for Chlamydiales. Finally, we highlight that approaches based on maximal exact matches may represent an advantageous compromise, achieving an intermediate level of computational efficiency while avoiding over-reliance on a single fixed *k*-mer length.

## 2 Introduction

The availability of assembled genomes of eukaryotes and prokaryotes has surged [17, 28], owing to advances in long-read sequencing technology [53] and assembly algorithms [43]. This new abundance of high-quality assemblies creates an ideal setting to reexamine the definitions and algorithms we use to study sequence similarity and evolution.

Robust metrics and models are needed to accurately quantify genome comparisons for different evolutionary rates across genomic regions and species. A basic model of DNA sequence evolution is Jukes-Cantor (JC), which assumes equal frequency of each base (or amino acid) and equiprobable substitutions [22]. More flexible models have also been suggested (e.g. K80, F81, GTR, PAM, BLOSUM) and applied [51]. These more complex models use scoring functions, assigning higher scores to substitutions that occur more frequently. In this way, alignments are driven toward identifying homologous segments descended from a common ancestor [2].

Average nucleotide identity (ANI) is a widely used measure for genomic similarity [21, 62, 63, 64]. ANI was originally proposed as a measure for delineating species and strains, specifically as an alternative to the labor-intensive DNA-DNA hybridization (DDH) technique [23, 40]. ANI has proven useful in other applications including building phylogeny [14] and guide trees [5, 40], improving the NCBI (National Center of Biotechnology Information) taxonomy [6], studying microbial presence in a metagenomic sample

Computing ANI is neither conceptually nor computationally trivial, despite the development of several ANI tools. What ANI truly represents is difficult to define because not only has the definition evolved over the years and holds different meanings across communities, but also definitions are tied to software tools, with their own scoring functions and complex heuristics.

In this review, we explore the state of the art in estimating sequence similarity and evolutionary distance between genome sequences. We propose an accuracy measure based on rank correlation with tree distance. Using this, we examine different ways in which researchers have defined the core notion of Average Nucleotide Identity (ANI). We then compare and contrast current approaches, highlighting methods on either end of the efficiency-versus-accuracy spectrum. Finally, we suggest how future methods could achieve even more favorable combinations of speed and accuracy. Our EvANI evaluation framework is available as an open source software tool at https://github.com/sinamajidian/EvANI.

### 2.1 Definition of ANI

The earliest uses of the term Average Nucleotide Identity (ANI) date to the 1990s when RNA polymerase gene sequences of Cyanobacteria [4, 39] were compared using the CLUSTAL multiple sequence aligner [15]. In 2005, a formal definition was introduced, with the aim of capturing similarity as measured using the molecular technique of DNA–DNA hybridization (DDH) [23]. In that work, predicted protein-coding sequences (CDS) from one genome (query) were searched against the genomic sequence of another genome (reference), and ANI was calculated based on conserved genes identified by BLAST (Basic Local Alignment Search Tool). The conserved genes were defined as BLAST matches with more than 60% overall sequence identity over an alignable region covering at least 70% of their nucleotide length in the reference.

Later, the meaning of ANI was widened to cover comparisons of whole-genome sequences [13]. To achieve this, the sequence of the query genome was divided into consecutive 1020-base fragments. This value corresponds to the size of the DNA fragments used in the DDH experiments. *In silico,* these 1020- base fragments were compared to the entire genomic sequence of the other genome (reference) in the pair using BLAST. The ANI between the query and the reference was calculated as the average sequence identity of all BLAST matches with greater than 30% sequence identity over alignable regions covering more than 70% of their length. They showed that a 95% ANI is equivalent to a DDH value of 70%, the threshold used for species delineation. In their study, they also performed a reverse search in which the reference genome is used as the query to provide reciprocal values. Their results show a small difference (less 0.1%) between the two reciprocal ANI values for each pair.

To improve computational efficiency, the JSpecies tool [44] used NUCmer (NUCleotide MUMmer) [9, 25] as a replacement for BLAST in order to compute the whole genome alignment (**Figure 1A**). In section 2.2, we will describe these in more detail. Recent methods define ANI simply based on a whole-genome alignment. Once a whole-genome alignment has been performed, ANI is computed as the fraction of aligned positions that are matches.

**Figure 1:**
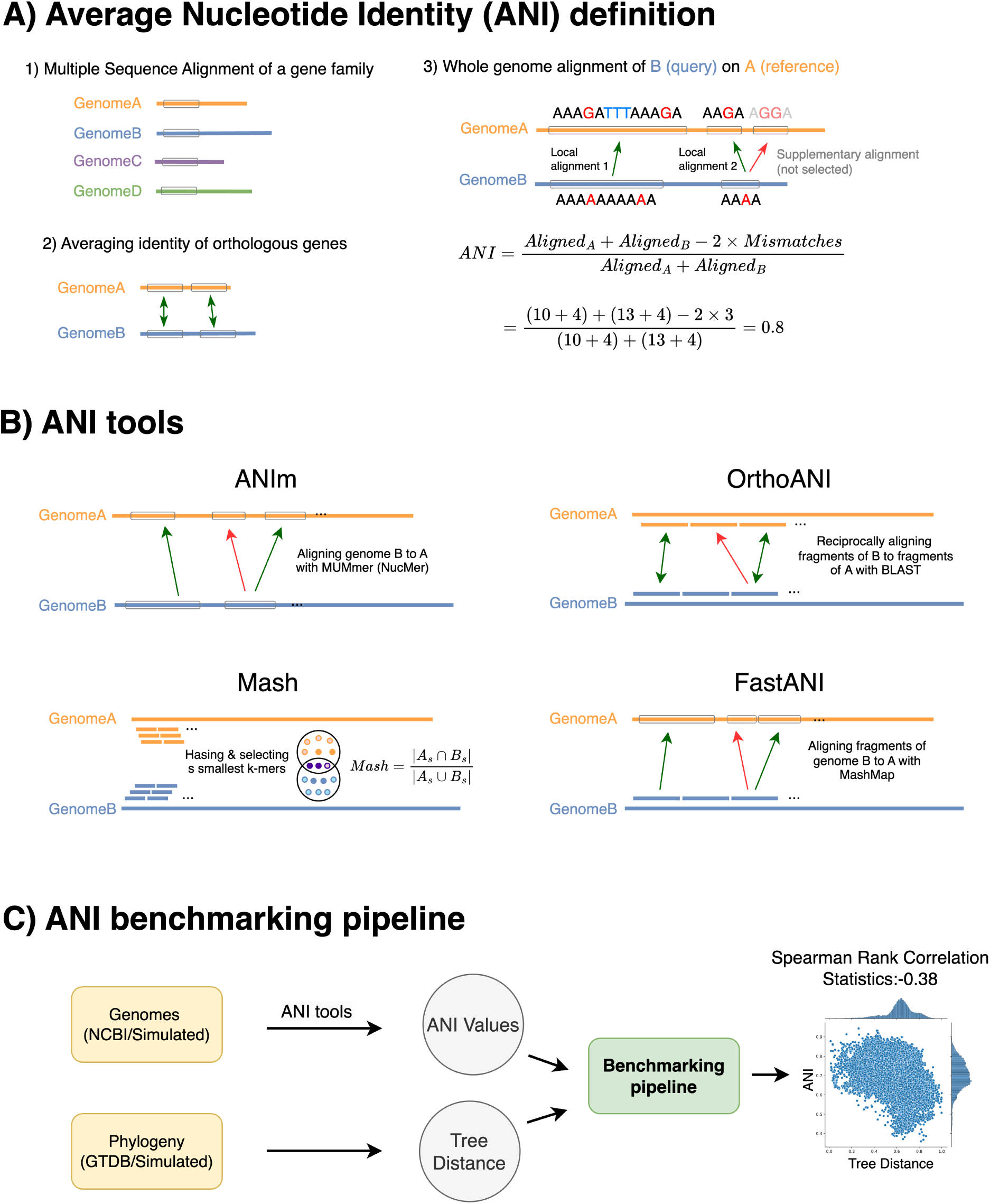
**A)** Average Nucleotide Identity (ANI) quantifies the similarity between two genomes. ANI is defined as the number of identical bases divided by the number of aligned bases. Historically, ANI was calculated using a single gene family for multiple sequence alignment. Another approach finds orthologous genes between two genomes and reports the average of similarity between their coding sequences. This method was later extended to whole-genome alignment by identifying local alignments and excluding supplementary alignments with lower similarity. **B)** Different ANI tools employ various approaches in calculating ANI values. ANIm, OrthoANI, and FastANI use aligners to identify homologous regions, whereas Mash uses k-mer hashing to estimate similarities. Only alignments with higher similarity represented by green arrows are included in ANI calculations, while red arrows, corresponding to paralogs, are excluded. **C)** The proposed benchmarking method evaluates the performance of different tools using both real and simulated data. It assumes that more distantly related species on the phylogenetic tree should have lower ANI similarities. This is measured by calculating statistics of Spearman rank correlation. We expect a negative correlation between ANI and the tree distance (scatter plot on the right).

While each of these definitions of ANI had practical motivations and interpretations, they did not yield the exact same definition. One definition is more concerned with how genes co-occur, while the other is more concerned with similarity of substrings that may or may not overlap genes. We also highlight some additional ambiguities in the definition and goal of ANI. First, some methods for computing ANI have the goal of computing average base-level identity over “alignable” regions only [13, 27, 44]. In these cases, the portions of one genome that fail to align to a counterpart in the other genome are totally excluded from the computation; that is, they are excluded from both the numerator and the denominator of the fraction computed. Whether or not this is a reasonable strategy depends on the scenario, as we will detail later. If the genomes being compared are distant, this requirement can result in an ANI estimate of zero or a value that is very close to zero. For instance, consider two scenarios: 1) two genomes with 90% ANI similarity over a 70% aligned region, and 2) two other genomes with 85% ANI similarity over a 100% aligned region. It is not clear which scenario represents the smaller evolutionary distance, even though the aligned region exceeds the threshold (70%). Furthermore, the threshold is often chosen arbitrarily,

One challenge in the definition of ANI lies in using a notion of conserved regions. The concept attempts to infer conservation through pairwise comparisons. However, the concept of conservation needs to be evaluated across several genomes in a pangenomics context. In other words, when identifying conserved regions—commonly referred to as core genomes—a pairwise approach is insufficient. Instead, comparisons across multiple genomes might be necessary, typically achieved through multiple genome alignment [59].

The primary approach intended in the ANI calculation is to capture regions of common ancestry using orthology [27]. When a distance metric aims to reflect a species tree, only orthologous regions should be taken into account. The rationale for including only orthologous regions is to avoid confounding effects from other evolutionary events, such as duplications. Consequently, duplicated regions should be excluded from the distance calculation to better conform to distances on the species tree. However, in practice, this goal is often not fully achieved, as identifying orthologous regions is challenging. Several methods for orthology inference have been proposed [1, 31] but they focus only on coding sequence regions. ANI tools often use reciprocal best hit as a proxy for finding orthologous regions. However, due to varying evolutionary rates across different genomic regions, the reciprocal best hits serve as a poor approximation [7]. It is also crucial to distinguish duplications that happened before or after speciation events, as this distinction impacts the orthology assignment.

Another limitation arises from using non-overlapping 1kb segments as the unit of comparison in the proposed definition. This approach interferes with finding the true borders of orthologs (or homologs). The proposed ANI definitions rely on alignment tools such as BLAST or MUMmer. Thus, calculation is at the mercy of heuristics used in the alignment process. BLAST, for example, employs a dynamic programming algorithm (i.e. local or global alignment) while MUMmer uses its own set of heuristics.

One important challenge is the prevalence of lateral gene transfer (LGT), *a.k.a,* horizontal gene transfer (HGT). This refers to the movement of genetic material between organisms that are not direct (vertical) descendants of each other. It could be argued that HGT has limited impact if the analysis is performed genome-wide, where ANI is calculated by averaging over all regions of similarity.

In conclusion, while the ANI concept is practically useful, it lacks a single, universally used definition. ANI’s reliance on alignable regions and pairwise comparisons introduces inaccuracies, particularly for distant genomes, as it excludes unaligned genomic portions and struggles with identifying conserved orthologous regions. Additionally, challenges arise from the use of fixed-width (e.g. 1kb) segments, from differing alignment heuristics, and from the impact of phenomena that have a more dramatic impact on both sequence similarity and tree shape, such as LGT.

### 2.2 Evaluation of ANI tools

We divide ANI-estimation tools into two categories: alignment-based and k-mer-based approaches. Alignment-based tools include OrthoANI, digital DDH, JSpecies, ANIb, and ANIm, with PyANI serving as a Python wrapper for the last two. K-mer-based approach include Jaccard and Mash (**Table 1**).

**Table 1:** A summary of the two broad categorizes of ANI tools including k-mer-based and alignment-based is provided along with their underlying software, links and references (see **Figure 1** for a visualization of how tools operate). Note that MUMmer refers specifically to its alignment subprogram, NUCmer. G2G: genome to genome, S2G: segment to genome, S2S: segment to segment.*:PyANI is a wrapper to ANIb/ANIm.

| Tools | Unit (default) | Alignment | Reciprocal | Underlying software | Software link | Ref |
| --- | --- | --- | --- | --- | --- | --- |
| Jaccard | k-mer | No | No | Dashing | github.com/dnbaker/dashing | [3] |
| Mash | k-mer (k=21) | No | No | MinHash | github.com/marbl/Mash | [38] |
| Dashing | k-mer (k=31) | No | No | Jaccard | github.com/dnbaker/dashing | [3] |
| ANIb | segment (1020bp) | S2G | No | BLAST | N/A | [13] |
| ANIm | genome | G2G | No | MUMmer | N/A | [44] |
| PyANI | * | * | * | BLAST/MUMmer | github.com/widdowquinn/pyani | [42] |
| JSpecies | genome | G2G | No | BLAST/MUMmer | jspecies.ribohost.com | [44, 45] |
| Digital DDH | genome | G2G | . | BLAST | ggdc.dsmz.de/ggdc.php | [33] |
| OrthoANI | segment (1020bp) | S2S | Yes | BLAST | help.ezbiocloud.net | [27] |
| FastANI | segment (3000bp) | S2G | Yes | MashMap | github.com/ParBLiSS/FastANI | [21] |

**Table 2:** The Wall-clock time of CPU time (in hours:minutes:seconds) of different ANI tools for simulated genomes (N=15; 105 genome pairs), Caldisericia (N=49; 1,176 genome pairs), and Cyanobacteriales (N=585; 170,820 genome pairs) using 48 CPU cores. K-mer approaches (Mash and Dashing Jaccard) are much faster than the alignment-based tools. ANIb did not finish for Cyanobacteriales in 24 hours.

| Tools | Simulated data |  | Caldisericia |  | Cyanobacteriales |  |
| --- | --- | --- | --- | --- | --- | --- |
|  | Wall-clock time | CPU time | Wall-clock time | CPU time | Wall-clock time | CPU time |
| Mash | 00:00:01 | 00:00:02 | 00:00:01 | 00:00:48 | 00:00:04 | 00:01:37 |
| Jaccard(Dashing) | 00:00:03 | 00:00:34 | 00:00:03 | 00:01:13 | 00:06:53 | 05:16:20 |
| FastANI | 00:00:10 | 00:04:14 | 00:00:07 | 00:03:37 | 01:02:47 | 33:40:50 |
| ANIm | 00:01:11 | 00:20:22 | 00:01:27 | 00:45:19 | 12:48:08 | 556:41:48 |
| ANIb | 00:03:02 | 00:41:47 | 00:03:57 | 00:57:37 | N/A | N/A |

#### The alignment approach

Early methods for ANI estimation focused on aligning genomes to each other. First, alignable portions of the two genomes were found with BLAST, as a tool for homology discovery. This was achieved by extracting consecutive blocks of 1020 bases of one genome, then aligning them to the other [13]. This was the approach taken for the method called “ANIb” [13, 44], which is implemented in the PyANI package [42].

A similar approach, called OrthoANI [27], divides both genomes into segments of 1020 bases. These segments are then aligned to each other to identify reciprocal best hits. ANI is calculated considering only the segments for which a reciprocal best hit was found. There are two notable differences between OrthoANI and ANIb (**Figure 1B**). First, only one genome is divided into windows in the ANIb method, whereas both genomes are divided into windows in the OrthoANI method. Secondly, in the ANIb analysis, the order of query and reference were changed to identify the conserved regions. However, a reciprocal best hit calculation is implemented in OrthoANI to find orthologous regions. The challenge with this approach lies in the arbitrary definition of homology boundaries (with blocks of 1020 bases), which may fail to reflect the true evolutionary history. The BLAST option used include -dust=no, -xdrop gap=150, -penalty=21, -reward=1 and -evalue= *e*^−15^. However, the rationale behind their selection of arguments of cost function or thresholds remains unclear.

Digital DDH is a web service that compute various distances based on local alignments (*a.k.a,* high-scoring segment pairs) identified with the BLAST aligner [33] (see **Table 1**). One distance measure is defined as 1 - a ratio, where the numerator of the ratio is the total length of all local alignments, including both alignments of reference vs query and query vs reference, and the denominator is the sum of both genomes’ lengths a.k.a alignment fraction (AF). Another distance measure is similar, but modifies the numerator by using the sum of identical base pairs. Besides these formulas, sampling schemes based on bootstrapping and jackknifing are used to calculate confidence intervals. Notably, the outputs of these two distance formulas can vary significantly. These distance values are converted to DDH estimates using generalized linear models from empirical reference datasets. For a test case of a simulated genome pair, the DDH estimates were reported as 100%, 71.3%, and 98.7%, corresponding to alignment length divided by total length (AF), identities divided by alignment length, and identities divided by total length, respectively. The authors recommend using the middle metric, reporting the lower similarity in this case. They argue that since the other two formulas use genome length in their denominator, thus these two are more prone to inaccuracies incurred by incomplete assemblies.

The ANIm method is based on the MUMmer whole-genome aligner [25, 44]. Some studies including HyperGen [61] have used ANIm as the “gold standard” method. ANIm reduces the high computational cost of BLAST-based methods, by replacing the core alignment algorithm with MUMmer’s fast maximal unique match (MUM) and maximal exact match (MEM) finding algorithms. MUMmer uses a 32-bit suffix tree data structure (up to version 3 [25]) or a 48-bit suffix array (in version 4 [32]) for fast match finding, and has flexible options for parallel processing. ANIm uses NUCmer, a wrapper program that uses invokes MUMmer and then uses local alignment to extend and combine matches found by MUMmer. NUCmer uses MUMs as anchors by default (MUM mode). MUMs are matches that are unique in both the reference and query. Alternatively, NUCmer’s maxmatch mode uses all anchor matches regardless of their uniqueness (i.e. MEMs). ANIm is available through the JSpecies online platform [45] and is also implemented in the PyANI package [42].

FastANI [21] is generally faster than both ANIm and ANIb, and works aligning segments of 3kb extracted from one genome to another using the MashMap aligner [20]. The process begins with indexing the genome, and subsequently finding all alignments using a a winnowed-MinHash estimator to measure similarity between the 3kb segment and regions of the other genome. This approach operates under the assumption that k-mers follow a Poisson distribution, which might not be the case in practice. Since identifying matching k-mers is a step within the MashMap aligner, FastANI could be considered either alignment-based or k-mer-based.

#### The k-mer approach

An alternative approach for calculating the distance between two genomes involves decomposing the genomes into their constituent k-mers and summarizing these in a “sketch” data structure. A sketch functions as an approximate version of a set data structure. Sketches can be queried to estimate set cardinalities, as well as the similarity between two sets as measured by the Jaccard index. The Jaccard index is defined as the ratio of the number of distinct shared k-mers between the two genomes divided by the total number of distinct k-mers in either genome:

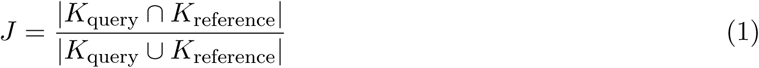

where *K_i_*is the set of all k-mers in the genome *i*, ignoring their multiplicity (i.e. each distinct k-mer counts once).

In their seminal study, Ondov *et. al.* described the Mash method, which uses the MinHash sketch to estimate pairwise distance. Instead of computing the Jaccard index overall all distinct k-mer, Mash computes it over a sketch, which is much smaller, approximate representation of the set. The MinHash sketch is constructed by hashing each k-mer from the input sequence and selecting the *s* smallest hash values. This reduced sketch enables fast, approximate set operations, such as intersection, facilitating efficient distance calculations:

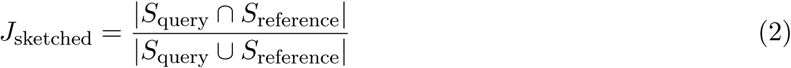

where *S_i_*’s is the set of hash values in the sketch built over genome *i*’s k-mers. The Mash index (similarity) is calculated as below

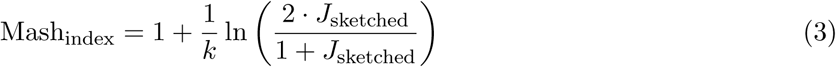

This calculation is based on the expectation that the number of mutations in a k-mer is *kd*, where *d* represents the mutation probability. Under a Poisson model, assuming unique k-mers and random, independent mutations, the probability of no mutation occurring in a k-mer is given by *e*^−^*^kd^* (for details, see page 10 of [38] and page 14 of [12]).

K-mer approaches like Mash are much faster than alignment-based approaches but typically achieve lower accuracy, as we discuss in Results. However, Mash is a widely used and successful tool, as are similar tools such as Sourmash [19] and Dashing [3], highlighting the utility of methods that sacrifice accuracy for computational efficiency.

A major goal of this review is to investigate specific reasons why *k*-mer based methods lose accuracy compared to alignment based methods. To this end, we will catalog their key assumptions and heuristics. Specifically, Mash employs a fixed k-mer length and applies a sketching step with a predetermined sketch size, limiting its focus to a sampled subset of k-mers. This *k*-mer-based strategy overlooks the collinearity of genomic segments and does not perform alignment at the base level, relying solely on exact k-mer matches. Once the statistics of shared k-mers are found, Mash calculates distance based on the assumption that k-mers are unique and independent of each other [12], ignoring the fact that two k-mers with a long shared suffix/prefix are more likely to co-occur than two k-mers with no shared suffix/prefix. Other k-mer-based tools include Bindash [55, 56] (which uses b-bit one-permutation rolling MinHash), HyperGen [61] (which incorporates hyperdimensional modeling of Dothash [37]), Gsearch [72], Skmer [48] (designed for raw reads), vclust [58] (designed for viral data), and skani [50] (for metagenomics which uses FracMinhash and k-mer chaining [18]).

## 3 Results

We developed a workflow to evaluate the performance of ANI tools under different scenarios and considering both distant and closely related species. We simulated genome evolution using the ALF simulator [8], varying amounts of duplication, lateral gene transfer (LGT), and mutation [8]. This simulation framework generates genomic sequences along with a phylogenetic tree that includes branch lengths, providing true distance values between genome pairs (**Figure 1C**). Additionally, we incorporated real genomes from NCBI RefSeq and phylogenetic data from both the NCBI taxonomy and GTDB [40, 41]. We worked from the premise that if an estimate of ANI is a good distance measure, it should accurately reflect the distances inherent in the species tree. In other words, a higher ANI value between two genomes should correspond to a smaller genetic distance on the tree, as indicated by a smaller sum of branch lengths between them.

### 3.1 Sketching of k-mers retains most of the essential information for distance calculation but there is no one optimal k-mer length across clades

Here, we aimed to evaluate how well a k-mer-based tool such as Mash and Jaccard can capture evolutionary distances. Tree distances represent the evolutionary divergence between genomes, providing a reference for assessing the accuracy of calculated distance measure. Thus, we investigated the correlation between the tree distance and the ANI estimates in different scenarios of genome evolution with varying degrees of genetic divergence.

Mash has two main parameters: k-mer length and sketch size. A larger k-mer length generally provides greater specificity, while smaller k-mers may offer more sensitivity [67]. However, large genomes might share small k-mers by chance, rather than due to sharing homologous regions.

The sketch size refers to the number of k-mers that are sampled and used in the analysis. A smaller sketch set enables fast comparison of genomes. The default parameters in Mash are k=21 and s=1000. We simulated the evolution of 15 genomes with varying mutation rates, from 5 to 100, resulting in average ANI values from 97% to 78%. The unit of mutation rate is point accepted mutation (PAM) and 100 PAM corresponds to one substitution per site on average (see the Methods section). The simulation outputs include the genomic content of species and the phylogenetic tree, which shows how samples are related to each other. The simulation experiment was repeated five times.

In our experiments with simulated data, we found that the default sketch size of 1000 was too small, leading to a Spearman correlation p-value of 10^−25^ for datasets with more distantly-related species (mutation rate of 100), whereas a p-value of 10^−151^ could be achieved using Mash without sketching (i.e. Jaccard). Increasing the sketch size from 1000 to 500,000 improved the Spearman p-value from 10^−25^ to 10^−73^ for mutation rate of 100. Varying the k-mer length for also affected distance estimation, with lengths of 15 or 17 yielding the best correlation (**Figure 2**).

**Figure 2:**
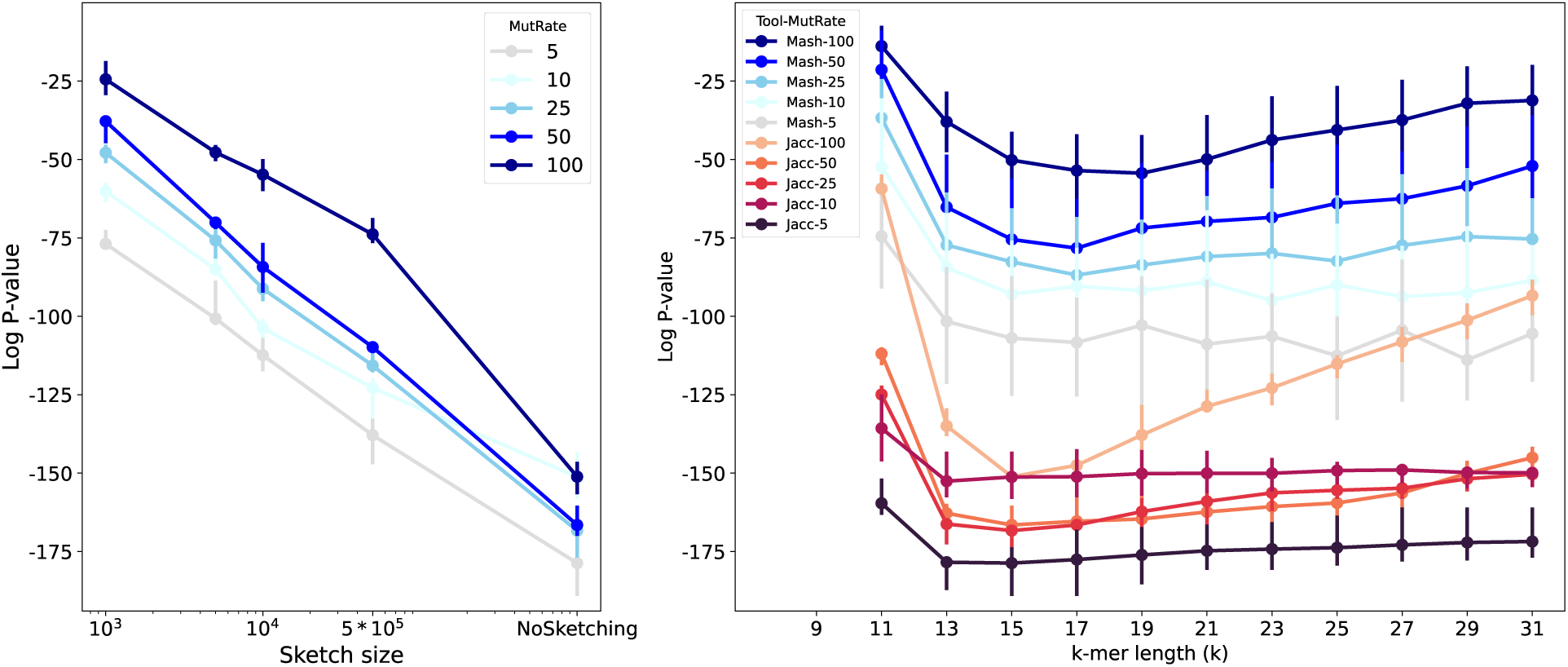
The *log*_10_ p-value of Spearman’s rank correlation test between ANI (calculated using Jaccard or Mash distance) with tree distance based on the true phylogenetic tree for simulated evolution of 15 genomes. (**Left**) Mash with k-mer length of *k* = 15, (**Right**) Mash with sketch size of *s* = 10000. The mutation rate for simulated genome evolution varies from 5 to 100. Overall, increasing the sketch size improves the rank correlation, decreasing the p-value. Also, the optimal k-value that minimizes the p-value is different for different scenarios.

We also analyzed real genome assemblies, focusing on six clades: *c_Caldisericia* (with *N* = 49 genomes), *o_Bacillales A* (*N* = 249), *p_Aquificota* (*N* = 132), *f_Neisseriaceae* (*N* = 145), *o_Cyanobacteriales* (*N* = 585), and *o_Chlamydiales* (*N* = 288). To calculate Spearman’s rank correlation, we considered the tree distance from the GTDB phylogenetic tree. The results are depicted in **Figure 3**. Notably, for Cyanobacteriales and Chlamydiales, statistics across different k-values revealed distinct minima at *k* = 11 and *k* = 20. No single value of k consistently optimized the rank correlation, highlighting a limitation of approaches that estimate evolutionary distance using fixed-length k-mers.

**Figure 3:**
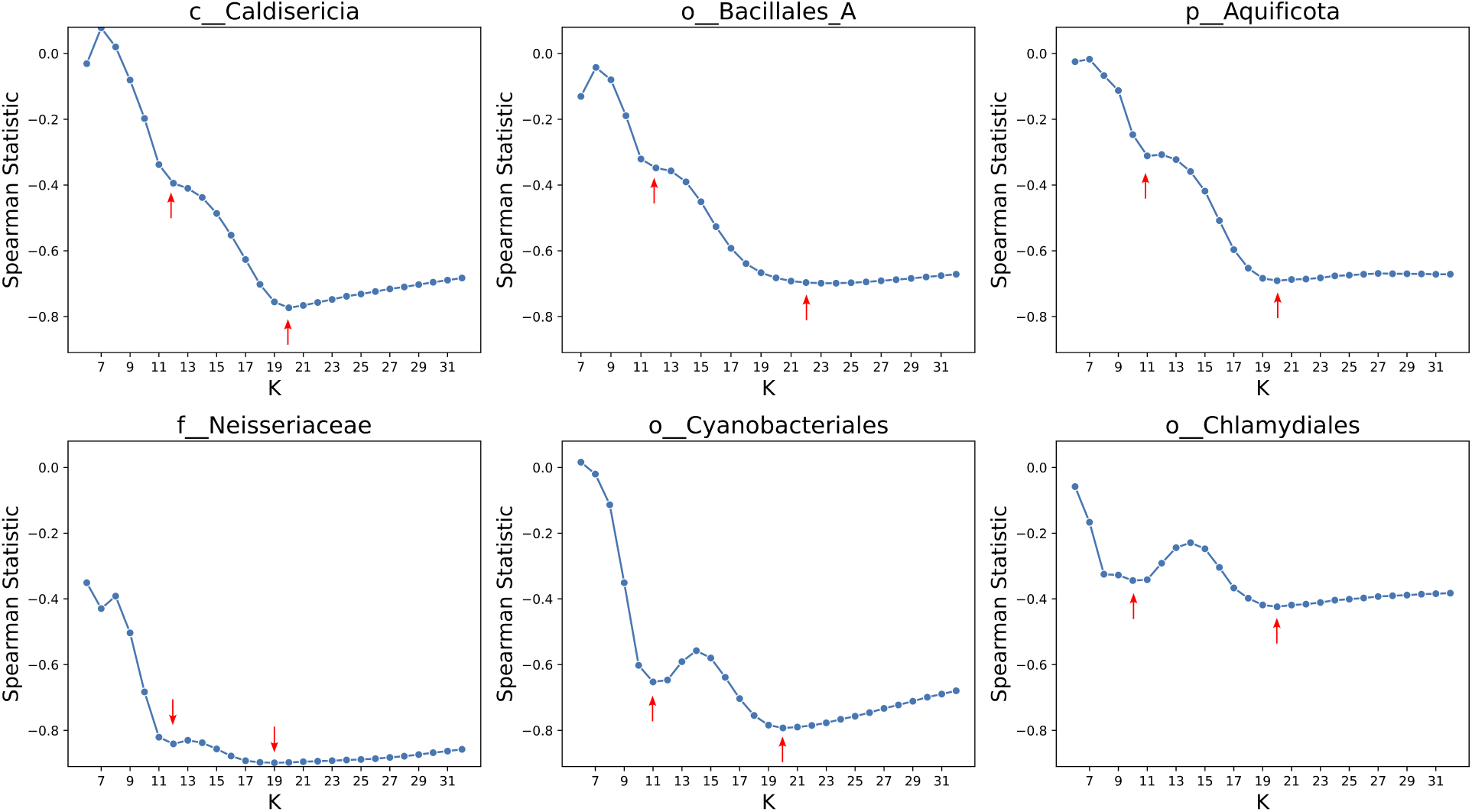
The statistics of the Spearman rank correlation test for six clades including *c_Caldisericia, o_Bacillales A, p_Aquificota, f_Neisseriaceae, o_Cyanobacteriales*, and *o_Chlamydiales*. Red arrows show a local minimum or a notable change in the statistics. Interestingly, the statistics for Cyanobacteriales and Chlamydiales across different k values showed two distinct minima. There is no single K value for the k-mer length that optimizes the statistics for estimating evolutionary distances. These lines of evidence highlight the fundamental limitation of k-mer-based approaches.

We hypothesized that the two local minima corresponded to k-mer lengths that contributed orthogonal information about evolutionary distance. If true, combining the information from both k-mer lengths should yield improved rank correlation. For the order Chlamydiales, the minima were at *k* = 10 and *k* = 19. We computed a merged ranking by averaging the ranks of each genome pair calculated using 10- mers and 19-mers, then re-ranking according to these averages. The distances derived from the merged ranks improved Spearman correlation (**Figure 4**), supporting our hypothesis. An analysis that used GC content specifically in place of the 10-mers did not improve rank correlation in the same way (see **Supplementary Figures 1-2**), indicating that integrating 10-mers does not simply add information about GC content.

**Figure 4:**
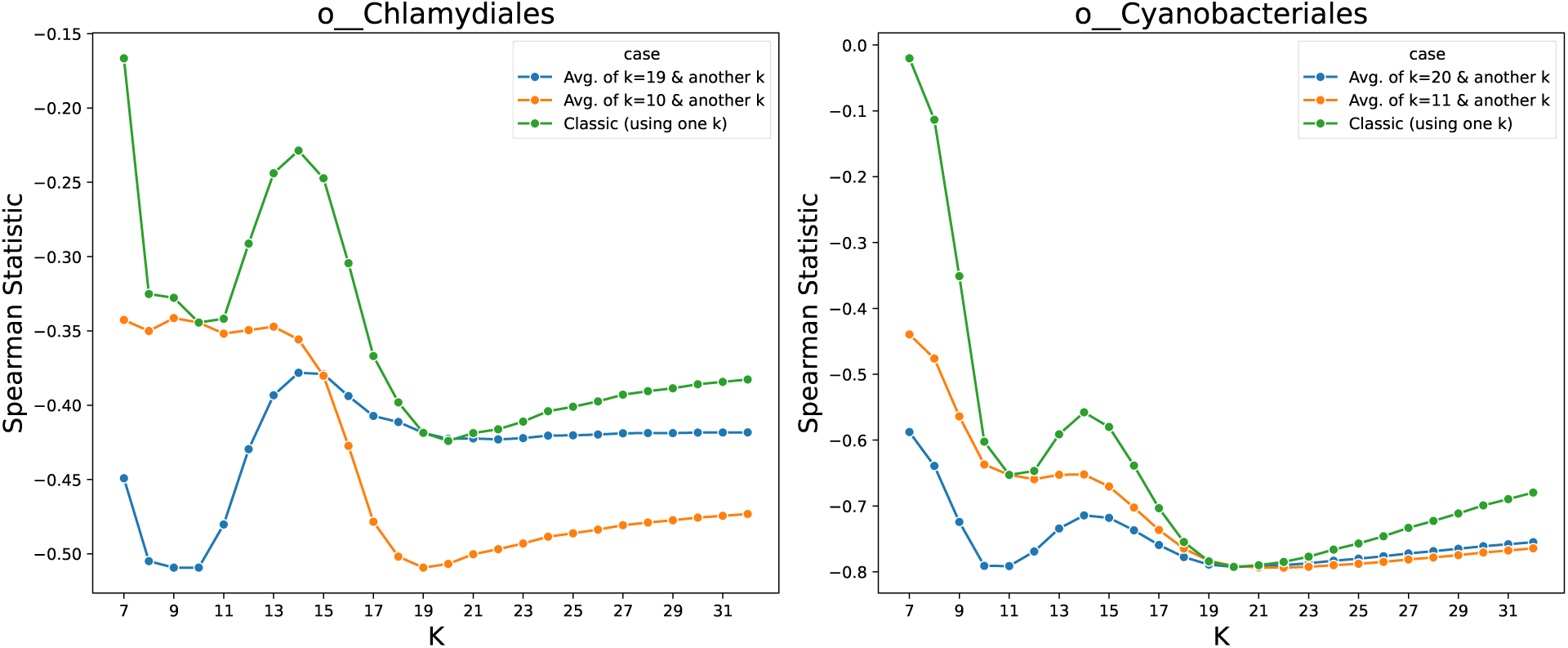
The Spearman rank correlation test between the distance on the GTDB tree and the Jaccard index. For the orders of Chlamydiales and Cyanobacteriales, two distinct k-values performed well (green). Using both sets of 10-mers and 19-mers (for Chlamydiales) to find distance ranks improves the statistics (blue/orange). This demonstrates that small and large k-mers can capture complementary information since using both resulted in a better estimation

### 3.2 Mash is robust to duplication and lateral gene transfer (LGT) while ANIm and FastANI are not as robust, but often perform better

We also conducted simulation experiments to study the impact of duplication and lateral gene transfer (LGT) on distance estimation. We used the ALF simulator [8] to generate genomes under duplication rates of 0.05%, 0.1%, and 0.2% or LGT rates of 0.01%, 0.05%, 0.1%, and 0.2%. We simulated each scenario five times and reported the median.

We ran Mash, FastANI and ANIm with different parameters. For Mash, we varied k-mer lengths and sketch sizes. For ANIm, we varied the minimum length requirement for the maximal unique matches (MUM). For FastANI, we varied the minimum fraction of genome shared (**Figure 5**) and the fragment length (**Supplementary Figure 4**). We observed that Mash was robust to increasing duplication rate but showed moderate sensitivity to higher LGT rates, with the impact becoming more pronounced for larger sketch sizes (see **Figure 5**). For example, for *k* = 15 and *s* = 50000, the averaged p-value for data without LGT was 10^−147^ but having LGT rate of 0.002 worsened this correlation to a p-value of 10^−106^. This matched our expectation that a higher rate of LGT adversely affects distance estimates since more genes can move among species in a manner that contradicts the species tree. The results also show that when the sketch set is large, the LGT has a more pronounced impact on Mash as mentioned earlier. On the other hand, for a sketch size of 1000, this value changes from 10^−75^ to 10^−69^, indicating some reduction in the accuracy of the estimate (see **Figure 5D**).

**Figure 5:**
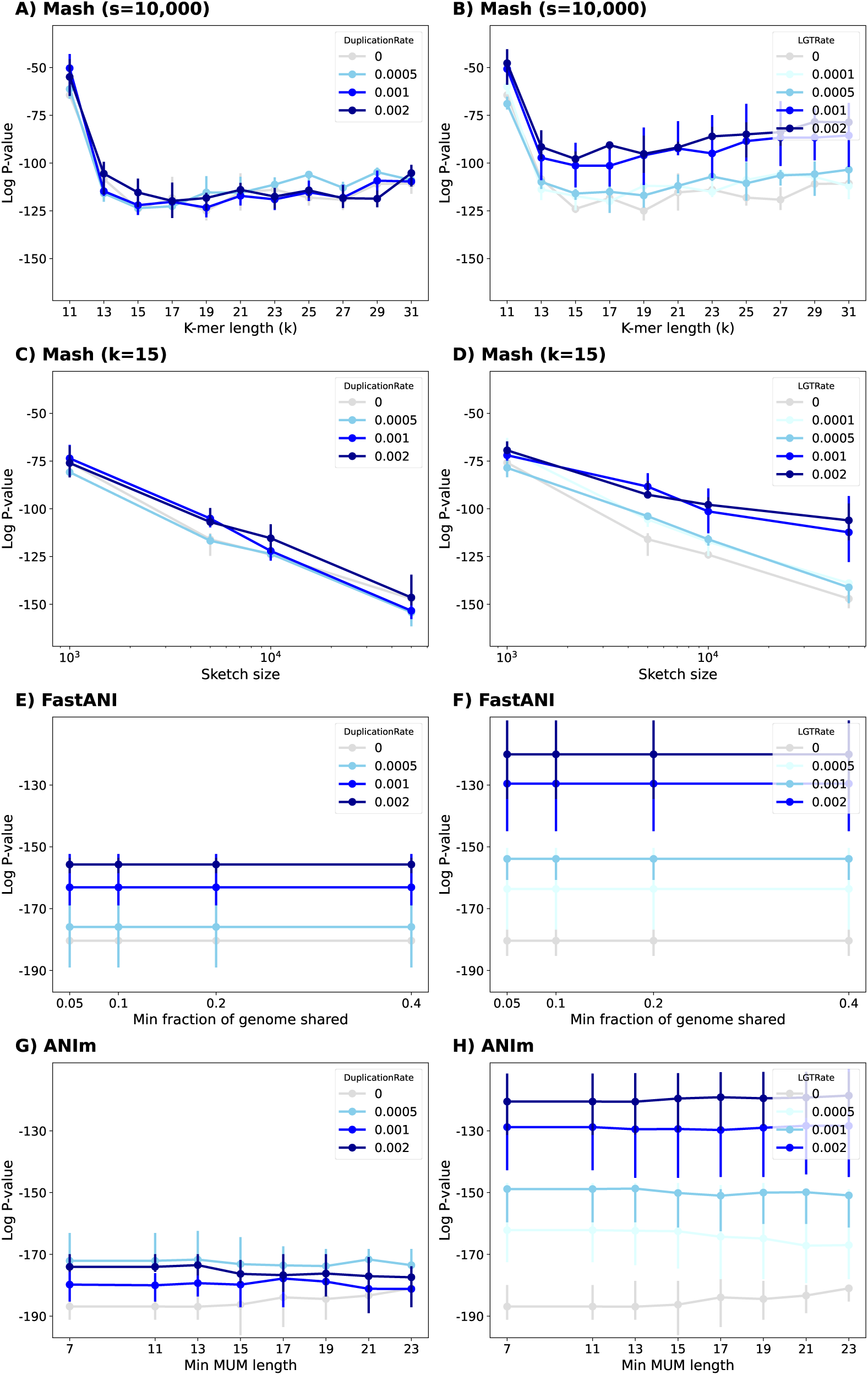
The ALF simulator was used to generate related genomes under two series of evolutionary scenarios. One series simulated duplication rates of 0.05%, 0.1%, and 0.2% (**left column**). Another series simulated lateral gene transfer (LGT) rates of 0.01%, 0.05%, 0.1%, and 0.2% (**right column**). **(A-D)** We ran Mash with different k-mer length and sketch size parameters. The alignment-based tools for estimating ANI includes FastANI (**E-F**) and ANIm (**G-H**). See **Supplementary** Figure 4 for the impact of fragment length on FastANI.

In contrast, alignment-based ANI estimation tools including FastANI and ANIm exhibited reduced performance as LGT and duplication rates increased. For these tools, the fragment length in FastANI or the minimum length of maximal unique matches (MUM) in ANIm did not significantly affect the Spearman correlation (**Figure 5E-F**).

So far, we studied the impact of duplication and LGT on the performance of each tool individually. Here, we want to draw a comparison among tools. Note that most tools have accuracies that fall into overlapping ranges, except Mash which has appreciably worse correlation than the rest (**Figure 6(right)**). By increasing the amount of LGT, the log Spearman p-value for all tools increases, i.e. their estimate exhibit worse correlation. The Jaccard index (found using Dashing in its `--use-full-khash-sets` mode) has accuracy similar to that of alignment-based tools like ANIb and ANIm for low LGT, which is surprising. Jaccard’s p-value changes drastically from 10^−186^ to 10^−107^ as the LGT rate increases to 0.002. Mash is less sensitive to LGT, achieving moderate p-value, changing from 10^−124^ to 10^−97^. This difference in behavior between Mash and Jaccard may be attributed to the Mash’s sketching. Given that the the number of exchanged elements due to LGT is limited, most of these elements are likely excluded from the sketched set of k-mers, resulting in minimal impact on the distance calculation by Mash. In contrast, Jaccard incorporates k-mers from the entire genome, including these exchanged elements, into the distance calculation, even though it should not. Since, these LGT genomic elements do not adhere to the vertical evolutionary history represented by the species tree, they should not be included in the distance calculation. Alignment-based approaches showed a decline in performance as the LGT rate increases, likely for the same reason.

**Figure 6:**
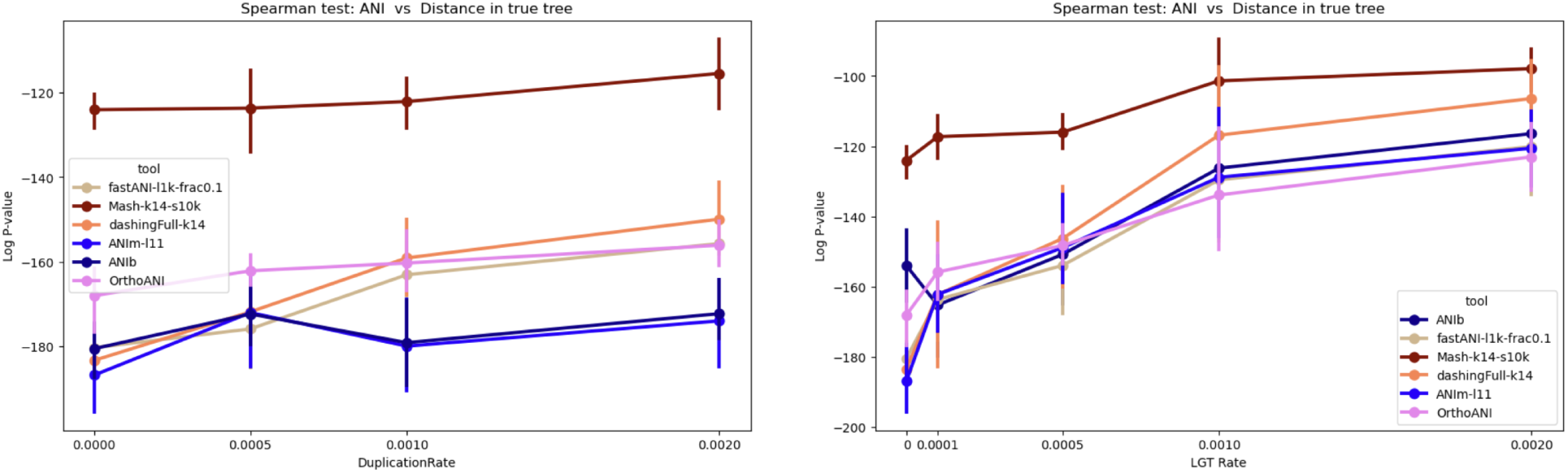
Comparing alignment-based tool and Mash on datasets with different LGT and duplication rates (summarizing the Figure 5). The alignment approaches perform better than k-mer-based Mash in Spearman correlation test. ANIm and ANIb are the best tools on the simulated data. (`fastANI-l1k-frac0.1`: FastANI with a fragment length of 1000 and minimum fraction shared genome of 0.1, `Mash-k14-s10k`: Mash with k-mer length of 14 and sketch size of 10,000. `ANIm-l11`:ANIm with a minimum MUM length of 11. `DashingFull-k14`: The Jaccard index was calculated with the Dashing tool which was run in the mode `--use-full-khash-sets`.

We also noted that in scenarios with high LGT or duplication, the ANIm and FastANI methods showed worse correlation. Specifically, ANIm’s p-value moves from 10^−186^ to 10^−120^ (10^−174^) as the LGT rate (duplication rate) increases from 0 to 0.002. Similarly, FastANI’s p-value moves from 10^−180^ to 10^−120^ (10^−155^) under the same conditions. But even in these cases, ANIm and FastANI exhibit better corelation compared to Mash (**Figure 5**).

The runtime of each tool varies drastically, chiefly depending on whether they are alignment-based or k-mer-based. Since the N=15 simulated genomes have a length of around 5 Mbp each, Mash and Jaccard take only a few seconds to run, whereas ANIm and ANIb need minutes to estimate ANIs between the full set of 105 genome pairs (=*N* × (*N* − 1)*/*2). This difference is more pronounced for the Cyanobacteriales clade, which compromising N=585 samples and so requires distances estimates for 170,820 genome pairs. ANIm took 13 hours using 48 CPU cores (556 CPU-hours) whereas FastANI needed 1 hour (34 CPU-hours). FastANI’s speed is likely due MashMap’s use of fast k-mer sketching methods. ANIb, which was run via PyANI on 48 CPU cores, did not complete this task for the Cyanobacteriales within a 24-hour time limit. Mash finished the task in less than two CPU minutes, again owing to its use of thanks to the fast sketching technique. Jaccard needed around 5 CPU-hours, which makes it six times faster than the fastest alignment-based tool, FastANI. While ANIm and ANIb provide more accurate distance estimates, especially for higher LGT and duplication rates in simulated data, they are considerably slower than the other methods. In summary, ANIb consistently provides the most accurate tree distance estimates but at a much higher runtime. FastANI’s speed is likely due MashMap’s use of fast k-mer sketching methods. ANIb, which was run via PyANI on 48 CPU cores, did not complete this task for the Cyanobacteriales within a 24-hour time limit. Mash finished the task in less than two CPU minutes, again owing to its use of a fast sketching method. The Dashing-based computation of the full-fidelity Jaccard index needed around 5 CPU-hours, making it six times faster than the fastest alignment-based tool, FastANI.

To study how different tools mimic the behavior of ANIb (or ANIm), we compare each tool versus ANIb (or ANIm) in terms of the rank correlation between the pairwise distances computed by the tool versus those computed by ANIb (ANIm). The results show that ANIb and ANIm produce similar rankings, whereas Mash is less similar to both ANIb and ANIm (**Figure 7**). When there is a high rate of duplication, FastANI is more similar to the Jaccard index as computed by Dashing than it is to the distances produced by ANIb/ANIm.

**Figure 7:**
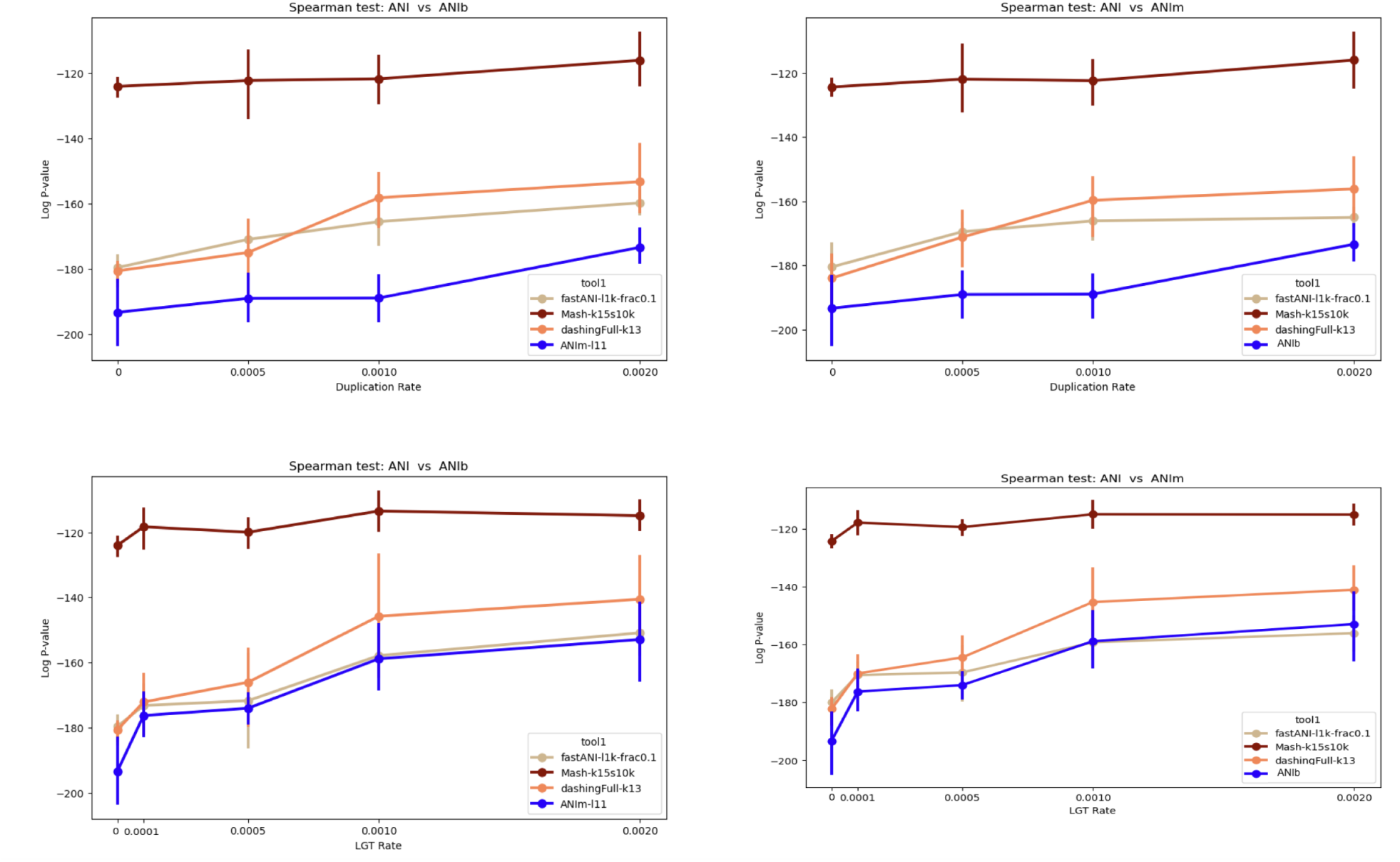
The *log*_10_ p-value of spearman correlation test between different ANI tools (FastANI, Mash, Dashing, ANIm) versus ANIb **(left)** using simulated data. We consider a range of LGT and duplication rates. **(right)** A similar analysis in comparison to ANIm. The results show the strongest rank correlation between ANIm and ANIb.

### 3.3 Consideration of match uniqueness, match length and alignment fraction boosts ANIm’s accuracy

ANIm has a tendency to report an ANI of zero for genomes pairs that are moderately distant. In a real dataset, we noticed that this was due to a lack of matches exceeding NUCmer’s minimum MUM length parameter. Note that so far and for the following analysis, NUCmer was run in MUM mode. Later we discuss the impact of using NUCmer in MUM mode compared to its maxmatch (MEM-based) mode.

When we ran ANIm with decreased value of the minimum MUM length parameter from 20 to 14. This caused ANIm to report non-zero distance values in more cases (**Supplementary Figures 3 and 12**), somewhat improving the Spearman correlation for its distance estimates. To further understand the advantage of MUM-based methods, we also explored the effect of weighting ANIm results by the fraction of the genomes that aligned. This was attempted in previous studies [14, 50] and we hypothesized that the adjustment could become more important as the genomes become more distant.

In an experiment with 84 Cyanobacteraia genomes, we tested this approach. We observed that for this dataset ANIm has a slight positive correlation with tree distance (Spearman statistics= 0.052 and p-value= 0.0017, **Figure 8**). This result is might be unexpected, as ANIm is a similarity metric and should not correlate positively with tree distance. This could be justified by previously reported limitation of ANIm in distance calculation when species are distant [50, 62]. Interestingly, when ANIm values were adjusted by multiplying them with the alignment fraction (AF), the negative correlation was observed, with a Spearman statistic of -0.286 and the noticeably low Spearman rank p-value of 7.3 ∗ 10^−67^.

**Figure 8:**
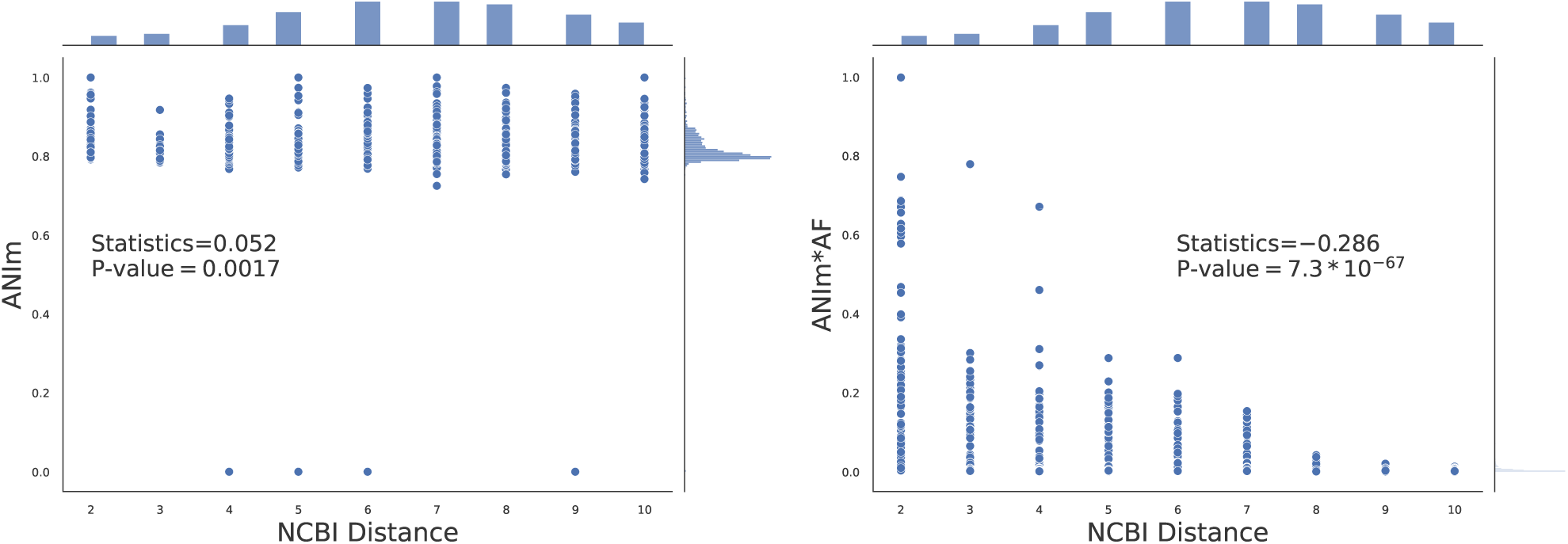
Impact of weighting ANIm with alignment fraction (AF) for distance calculation in Cyanobacteraia. Here, 84 genomes are considered for studying the correlation between ANIm (or *ANIm* ∗ *AF* ) versus tree distance from NCBI. Each point corresponds to a pair of genomes (See **Supplementary** Figure 10 when GTDB is used).

The simulation results showed that *ANIm* ∗ *AF* performs slightly worse than ANIm for closely-related species, but outperforms ANIm for more distantly related genomes. Under conditions where the mutation rate is 100 (producing divergent genomes in the evolutionary scenario), weighting improves the p-value from 10^−105^ to 10^−166^ (minimum MUM length=11, top left, **Figure 9**). When the minimum MUM length is set to 21 (default), the improvement is even more pronounced, from 10^−16^ to 10^−83^. For the same mutation rate (=100), the p-value of ANIb is 10^−176^. Interestingly, *ANIm* ∗ *AF* ’s performance surpasses ANIb’s when the mutation rate is 200 (**Supplementary Figure 11**).

**Figure 9:**
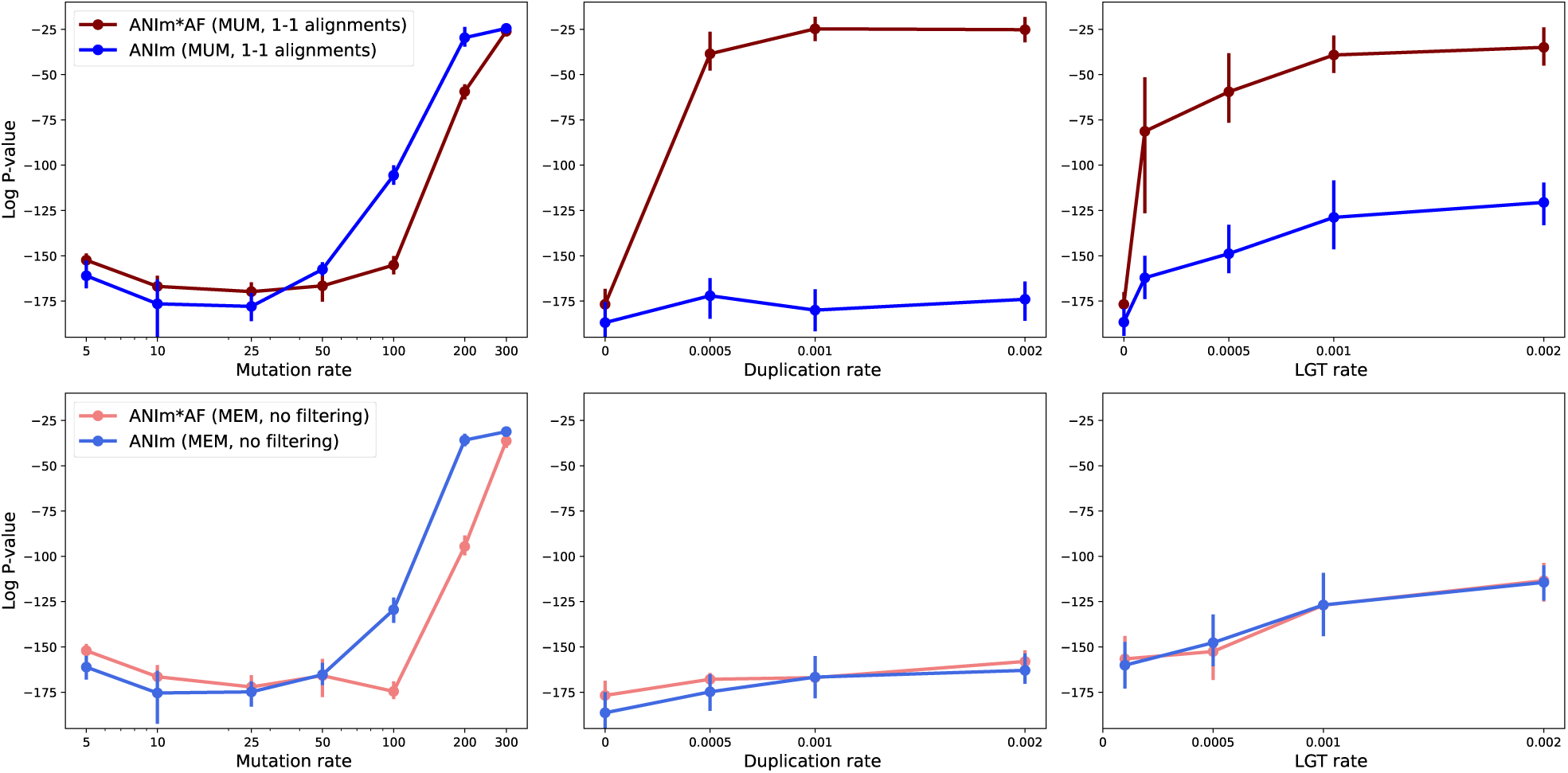
The Spearman correlation between ANIm (or ANIm∗AF) and tree distance for different mutation, duplication, LGT rates. The top row uses MUMs in alignment with MUMmer, followed by keeping only 1-to-1 alignments. The bottom row corresponds to using all maximal matches without any filtering. Higher mutation rates in the genome evolution result in more distant genome. Different minimum MUM length used in ANIm did not impact the result (**Supplementary** Figure 6). AF=alignment fraction.

On the other hand, our simulations also showed that the improvement achieved by using AF-weighted ANIm does not hold when there is a high rate of duplication or LGT (**Figure 9 (top row)**). Notably, we observed that the alignment fraction (AF, calculated with NUCmer in MUM mode) of genomes with duplication or LGT showed a much lower Spearman correlation with tree distance; specifically, the Spearman p-values increased from 10^−80^ (when no duplication/LGT exists) to 10^−17^ for a duplication rate of 0.002 and to 10^−7^ for an LGT rate of 0.002 (**Supplementary Figure 7**). Note that AF values are based on the 1-to-1 alignments that are found by filtering all NUCmer alignments. The fraction of unfiltered alignments correlates better with the distance (the last row of **Supplementary Figure 7**). This likely occurs because filtering removes a substantial portion of alignments in the presence of high duplication, which reduces the AF numerator (total alignment length) without changing the denominator (genome length). This leads to an alignment fraction that is lower than the true proportion of homologous regions.

The described analyses of ANIm were based on NUCmer’s MUM mode. In this mode, NUCmer uses MUMs as anchors which are maximal unique matches. Thus, NUCmer does not find all the available alignments due to the uniqueness of anchors. This is observed in a decrease in fraction of alignment to around 0.90 when duplicated genes are present (**Supplementary Figure 7-8**). To address this issue, we ran NUCmer in maxmatch mode which uses all maximal matches as anchors. Interestingly, *ANIm* ∗ *AF* performs similarly to ANIm even when high duplication or LGT is present (the bottom row of **Figure 9**). We hypothesize that since uniqueness of MUMs result in discarding some of the duplicated regions, creating big discontinuities and poor rank correlation. Consequently, *ANIm* ∗ *AF* does not appear to be a universally suitable measure for distance estimation. Developing an optimal weighting scheme that accounts for diverse evolutionary scenario including LGT and duplications, with their distinct alignment profile, could be a direction for future research. Overall, ANIm demonstrated superior computational efficiency compared to ANIb. Tuning match parameters (uniqueness and length) combined with AF weighing extends ANIm’s applicability to more distantly related genomes, achieving an accuracy comparable to that of ANIb.

### 3.4 Orthologous genes showed a stronger rank correlation with the GTDB true tree distance compared to the whole genome using k-mers

We investigated the impact of different genomic regions in distance calculation using k-mers. In an experiment on 44 genomes of *o_Bacillales A2*, we infer orthologous genes with the FastOMA tool [31]. We considered four types including the whole genome, all coding sequences (CDS), 100 random CDS genes, and 100 orthologous genes. We calculated the Jaccard similarity using each of the mentioned genomic regions separately. A typical k-mer-based distance method considers the whole genome. The results indicate that orthologous genes show a stronger correlation with species tree distances compared to whole-genome data for the clade of *o_Bacillales A2*. This is because these genes are responsible for speciation events, shaping the species tree. However, it is important to note that changes in coding sequences (CDS) alone may not provide sufficient resolution for calculating distances between subspecies or strains within a single species. Additionally, a limitation of this approach is its reliance on accurate orthology information of genes. Of note, gene annotation itself is a herculean task [46], impacting the orthology assignment [26], ultimately the distance estimation. This highlights the important point that methods should clearly specify their range of evolutionary distances they are designed to address. It also emphasizes the need for future tools with ability to design models that can integrate the strengths of approaches specialized in both short evolutionary distances and those tailored for longer evolutionary distances.

## 4 Discussion and Conclusion

A goal of many modern Bioinformatics methods is to efficiently compute values, e.g. Mash distance, that serve as proxies for evolutionary distance. In this study, we surveyed different definitions for average nucleotide identity (ANI) and examined how ANI is influenced by the heuristics employed in different ANI calculators. To achieve this, we developed a benchmarking workflow called EvANI to evaluate the performance of these tools across a diverse range of scenarios.

The term ANI has been used so broadly over the past years in the literature and it can refer to related concepts with different assumptions. We argue that the initial definition of ANI emerged as a practical approach, rather than a comprehensive definition. In the future, it is important to clearly define the assumptions underlying the use of ANI. We recommend being more explicit about the evolutionary phenomena that a study aims to measure, as these assumptions directly influence how a method calculates the ANI value, specifically how unaligned and duplicated regions are treated is of great importance.

As we discussed in this work, ANIb demonstrates impressive performance by aligning as much of the alignable sequence as possible, with unaligned regions likely reflecting a lack of homology. On the other hand, the k-mer approach takes the whole genome into consideration but underlying assumptions on k-mer length hinder accuracy to the level of ANIb. In other words, ANIb appears to be the most effective at capturing tree-based distances. However, its computational demands make it impractical for large-scale studies. Additionally, the scoring function used in BLAST (as the underlying software in ANIb) introduces assumptions that have not been thoroughly investigated. The current literature lacks an exploration of optimal scoring functions for alignment approaches in this context. Compared to k-mer-based methods, the space of possible scoring functions is much larger, indicating significant opportunities for further work. Designing scoring functions tailored to different genomic regions (e.g. genic vs. intergenic regions) based on their evolutionary histories might be particularly beneficial. Revisiting this problem will be crucial for improving ANI tools.

Overall, benchmarking the ANI is hard since there is no mathematical definition but only a practical description for ANI. In this study, we designed an approach that enables benchmarking ANI tools using distance on the tree. We highlighted different situations where different assumptions in the ANI tools played role and made a difference. This resulted in the recommendation that assumptions and limitations should be explicitly described and carefully considered when using these ANI tools.

The study demonstrates a direction for future research. Exploring alternative alignment tools, such as Minimap2 [29] or LastZ [49], and evaluating their performances could provide valuable insights. In the context of k-mer-based methods, investigating other sketching techniques such as minimizers [35] can enhance the performance. It also remains an open question as to whether we could achieve higher accuracy by combining different k-mer lengths together with spaced k-mers [30, 34], or more generally matching statistics [69]. Other lines of research could be on considering haplotypes variations of diploid combination [74] and large language models [73]. Finally, a comprehensive benchmarking of phylogenies [57] inferred from evolutionary distances, compared to those derived from gene marker-based [10, 11] or reconciliation-based approaches [54], would be highly valuable for the field.

## 5 Methods

The proposed benchmarking method requires two key inputs: the genomes and the phylogeny (**Figure 1C**). To evaluate how effectively different ANI tools measure evolutionary distance, we use a Spearman rank correlation test to quantify the correlation. The results are reported as correlation statistics or p-values in `log_10`.

### Real data

We utilized the phylogenetic tree from the GTDB resource [40, 41]. This phylogeny is constructed using multiple sequence alignments of 120 gene markers and thus includes branch lengths. To calculate tree distances, we employed the ete3 package [16]. We selected six different clades including *c_Caldisericia*, *o_Bacillales A*, *p_Aquificota*, *f_Neisseriaceae*, *o_Cyanobacteriales*, and *o_Chlamydiales*. For each clade, we retrieved the taxonomic IDs from GTDB and obtained their genomes from the NCBI Assembly database using command line `esearch -db assembly -query tax id | esummary | xtract -pattern DocumentSummary -element FtpPath GenBank`. We also used the phylogeny from the NCBI taxonomy. However, this phylogeny does not have branch lengths. Thus, we used topological tree distances (as integer values) in our method.

### Simulated data

We utilized the Artificial Life Framework (ALF) simulator [8] to model genome evolution via its command-line tool. For each scenario, 15 genomes were generated. We repeated this five times, and averaged the results across these replicates. Prior to genome simulation, ALF generates a species tree (phylogeny) by fixing speciation events (**Supplementary Figure 9**). The tree is sampled using a birth–death process, with a birth rate of 0.01 and a death rate of 0.001. The root genome consists of 100 genes, each with a length of 50 kb, resulting in genomes with an average size of 5 Mbp.

At each speciation event (an internal node in the species tree), two new species are generated, each inheriting the ancestral genome. These offspring genomes evolve independently, undergoing different mutations and accumulating distinct differences as the simulation progresses. We benefited from the point accepted mutation substitution model at the amino-acid level. The mutation rate influences the branch lengths on the tree, reflecting the extent of changes between an ancestral species and its descendants. Mutation rate is measured in point accepted mutation (PAM) and 100 PAM is equal to one substitution per site on average. Additionally, insertions and deletions (indels) occur independently of substitutions at a rate of 0.0001, following the ZIPF model [8].

Gene duplications occur randomly at the sequence level along with evolutionary events. In our simulation, we consider duplication rates of 0, 0.0005, 0.001, and 0.002 [8, 52]. Another event included in the simulation is lateral gene transfer (LGT), which allows a genome to acquire new genes. ALF randomly selects donor and recipient genomes as well as the genes to be transferred. The transferred genes are inserted at random positions in the recipient genome. We consider the LGT rate, varying from 0 to 0.0001, 0.001, and a maximum of 0.002 [8, 52].

### ANI Tools

We ran FastANI [21] version v1.34 using `fastANI --ql fastalist --rl fastalist -o distances.tab -- threads 48 --minFraction 0.1 --fragLen 3000` where `fasta list` is the list of input FASTAs. We considered a range of values for the minimum fraction of the genome that is shared (minFraction) and the length of the fragment (fragLen). Furthermore, we use OrtoANI [27] version v1.40 with `java -jar OAT cmd.jar -blastplus dir ncbi-blast-2.15.0+bin -num threads 48 -fasta1 ref -fasta2 query ref query.out`. This calculates the distance between two genomes, namely, fasta1 and fasta2. We feed OrthoANI with the executable of BLAST version 2.15.0.

We used PyANI [42] version 0.2.12 for measuring runtimes of ANIb and ANIm and `ran average nucleotide id -m ANIm -i fasta_folder -o out -v -l out.log --workers 48`, where `fasta_folder` is the folder including FASTA files.

PyANI [42] does not allow modifying options of the underlying aligner (ANIm or ANIb) and it soon ran out of memory with ANIb. We separately executed ANIb’s and ANIm’s underlying tools, which are NUCmer and BLAST, respectively.

Subsequently, we used Python functions from PyANI to compute the distance values. This approach allowed us to modify the minimum MUM length in NUCmer when we ran it with `nucmer -l 20 --mum -p ref query.out ref.fa` query.fa where -l is the parameter of minimum maximal uniq matches (MUMs). We also ran nucmer `--maxmatch` to examine the impact of match uniqueness on weighting ANIm with alignment fraction. The output of this step is alignment in delta format. Then, we filtered these delta files to select the best hit using `delta-filter -1 ref query.out.delta > ref_query.filtered.delta`.

To compute the Jaccard index, we used the Dashing tool [3] version v1.0.2-4-g0635 for different k-mer lengths, using `dashing dist --use-full-khash-sets -k 21 -p 48 -O distance.tab -- full-tsv fasta/*fa`. We executed Mash [38] version 2.3 in two steps of sketching and distance calculation. We ran `mash sketch -p 48 -o all -s 1000 -k 21 fasta/*fa` for creating sketches and mash `dist -p 48 all.msh all.msh -t > distances.tab` for creating the distance matrix. We varied the sketch size and the k-mer length with -s and -k arguments, respectively.

We ran FastOMA [31] version v0.2.0 to infer orthologous genes using `nextflow run FastOMA.nf --input folder proteomes --output_folder output` when the translated CDS genes were downloaded from NCBI.

We surveyed a wide range of methods for estimation of evolutionary distances. We developed the EvANI benchmarking framework and datasets to evaluate the accuracy of distance estimation algorithms. Bi-k-mer spectra provide better evolutionary distance estimates for Chlamydiales than a single k-mer.

## Data and Code Availability

The benchmarking workflow is available at https://github.com/sinamajidian/EvANI and benchmarking datasets are available at https://zenodo.org/records/14579845.

## Author contributions statement

SM drafted the manuscript and ran the experiments. BL and SM designed the experiments. BL, SH, MZ contributed to the analysis and revised the manuscript. All authors read and approved the final manuscript.

## Acknowledgments

We would like to thank Vikram Shivakumar for feedback and helpful discussions. S.M and B.L. were supported by NIH grants R35GM139602 to B.L.

## Key Points

- We surveyed a wide range of methods for estimation of evolutionary distances.
- We developed the EvANI benchmarking framework and datasets to evaluate the accuracy of distance estimation algorithms.
- Bi-k-mer spectra provide better evolutionary distance estimates for Chlamydiales than a single k-mer.
- BLAST-based ANIb effectively captures tree-based distances but is computationally demanding.

## 6 Supplementary Figures

**Supplementary Figure 1:**
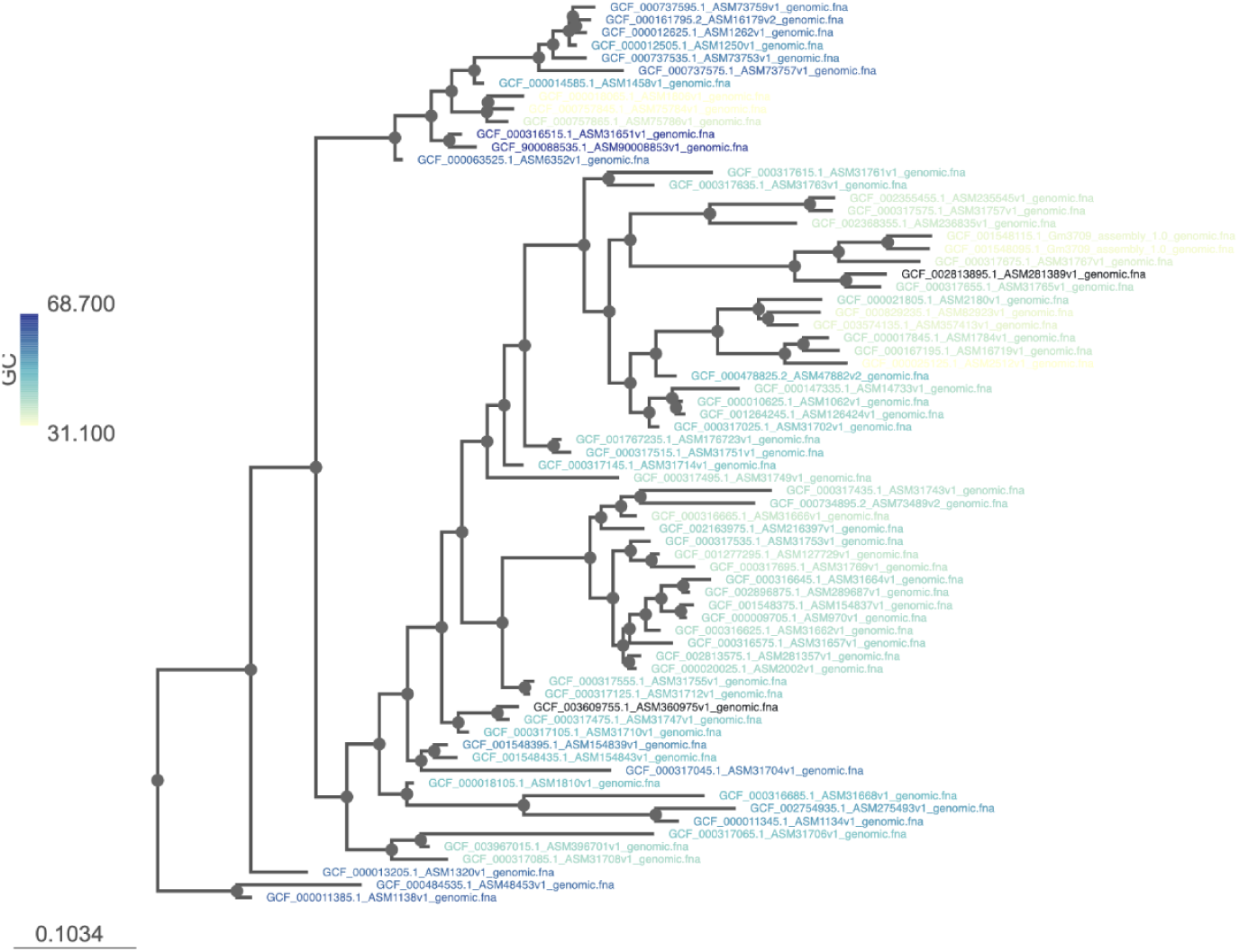
The GC content of species in Cyanobacteriota. There is high variation in GC content in Cyanobacteriota, ranging from 31% to 68%.

**Supplementary Figure 2:**
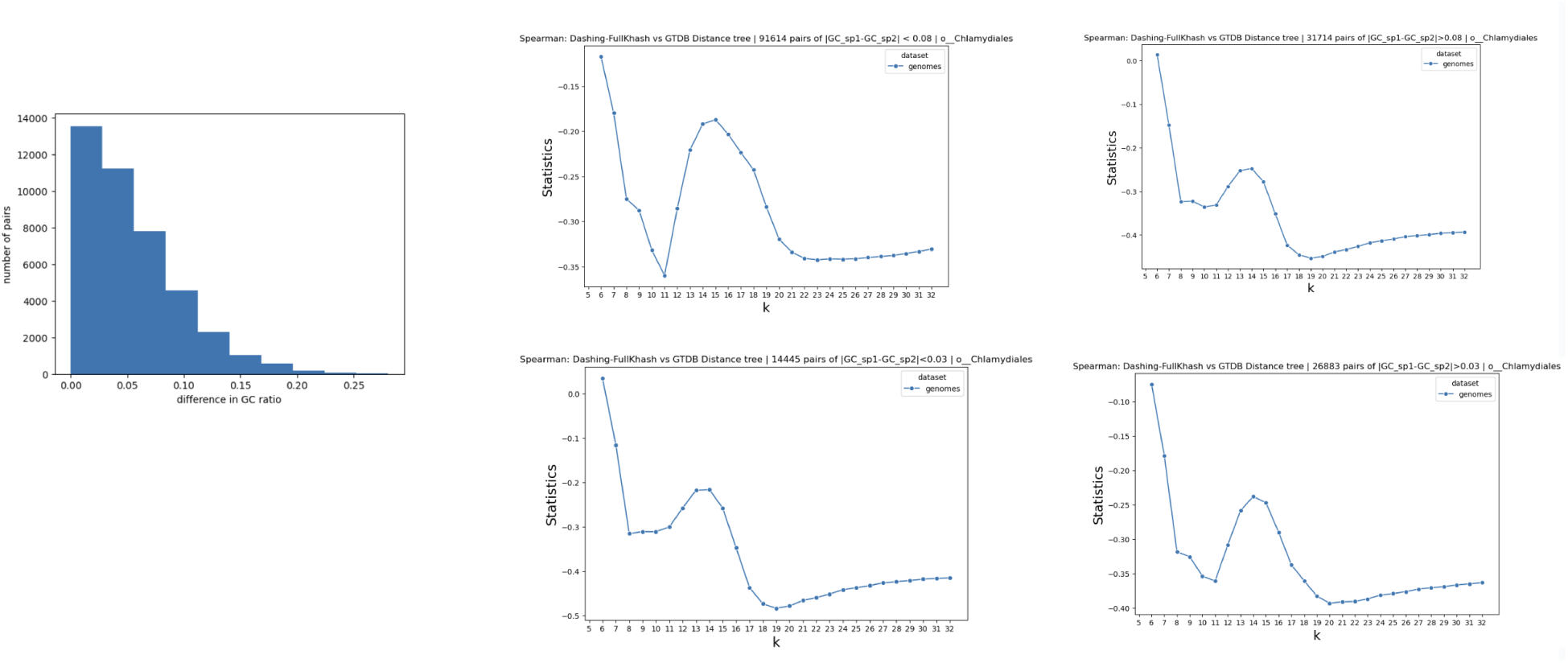
(left) The histogram of difference in GC ratio between every species pair. (right) In each subplot, we keep only a portion of pairs. If the GC difference had been the cause (of two minima separate considering all pairs), the sub-figure for pairs with similar GC would not have had two minima. Thus, difference in GC content does not justify the two local minima seen in Figure 3.

**Supplementary Figure 3:**
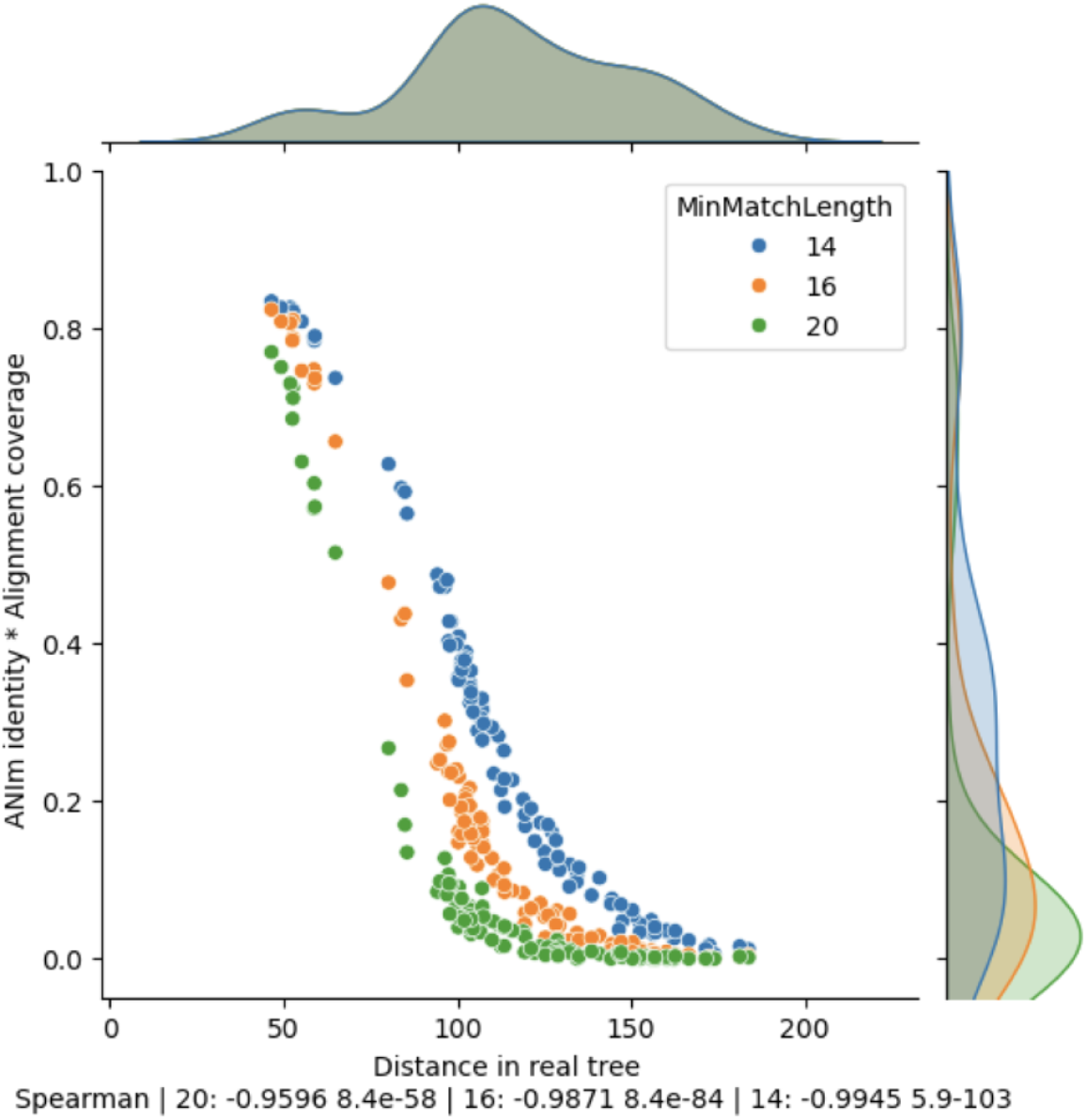
Impact of changing the MUMmer parameter minimum match length in distance estimation.

**Supplementary Figure 4:**
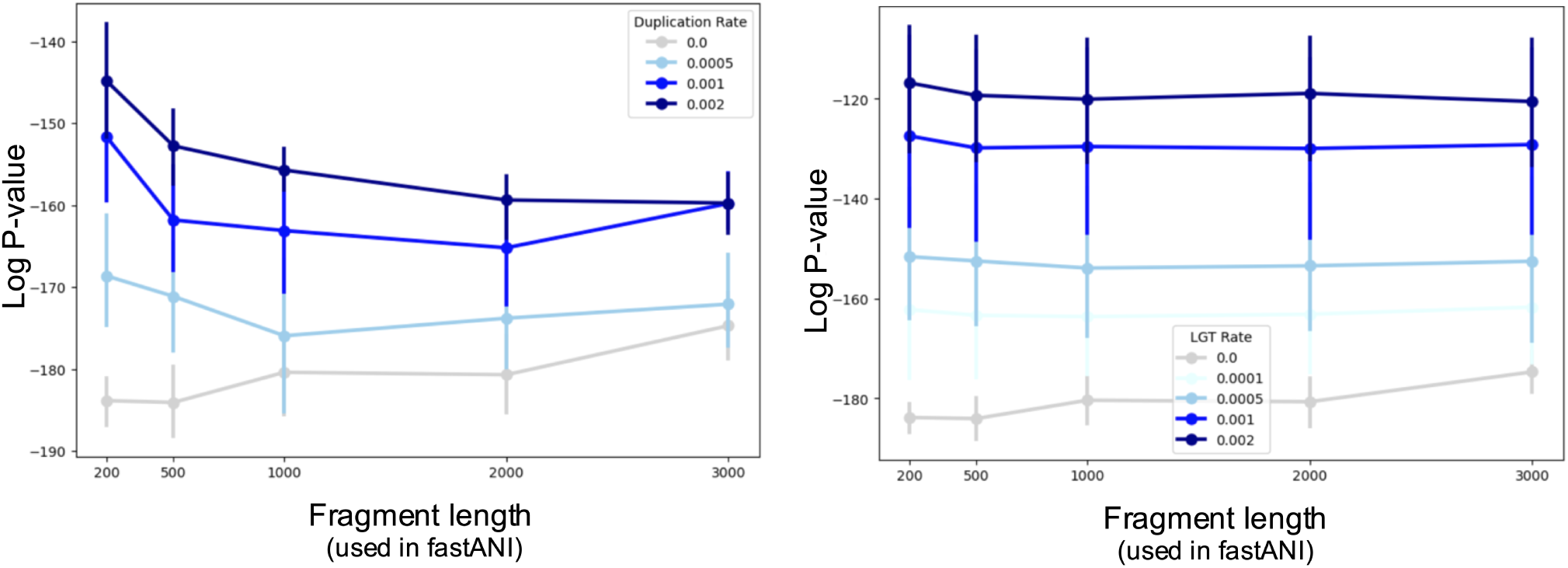
Impact of changing fragment length used in FastANI across different simulated data varying duplication and LGT rates.

**Supplementary Figure 5:**
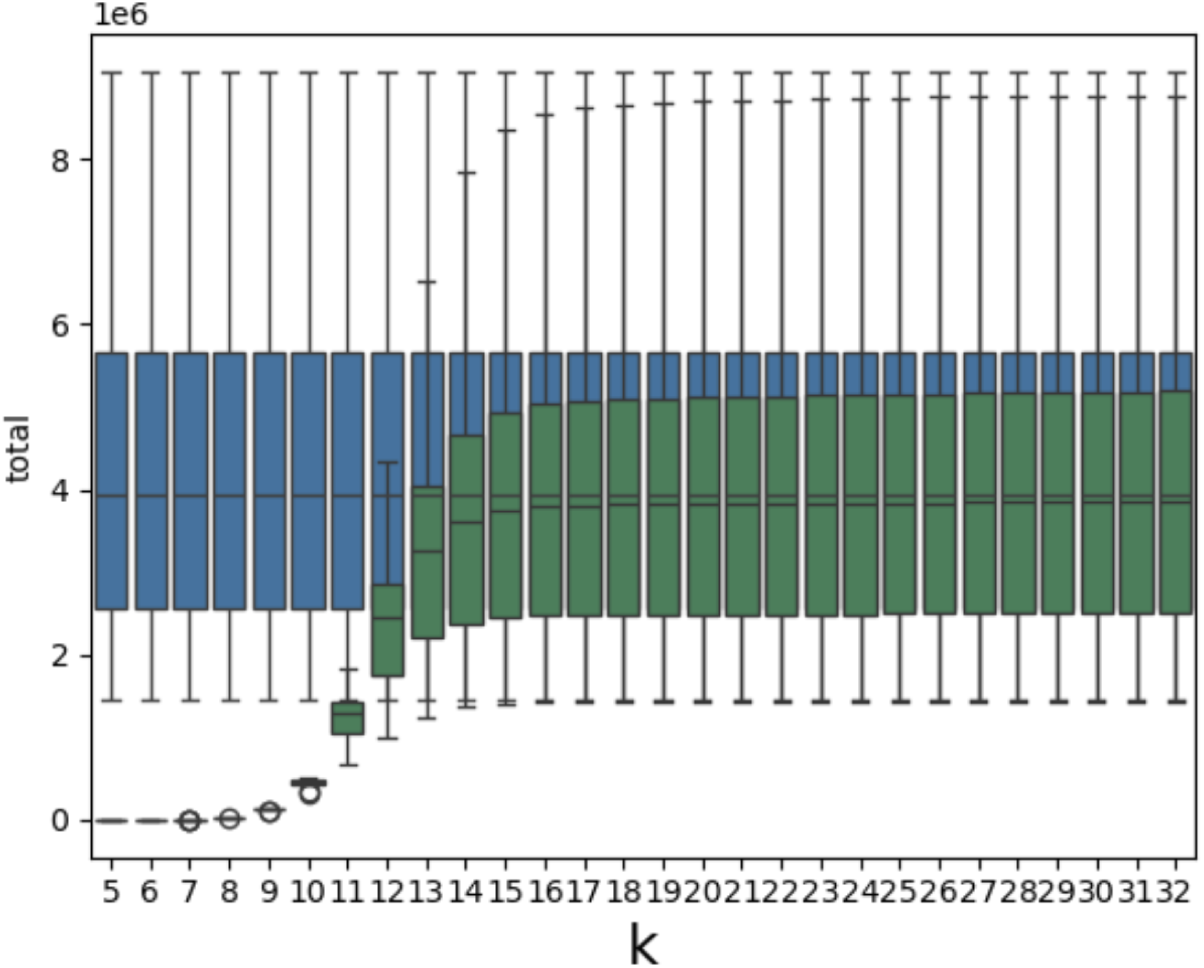
(blue) total number of k-mers, which is (almost) equal to the genome size, (green): number of unique k-mers found by the KMC tool. We can see there is a smooth increase in the number of unique k-mers.

**Supplementary Figure 6:**
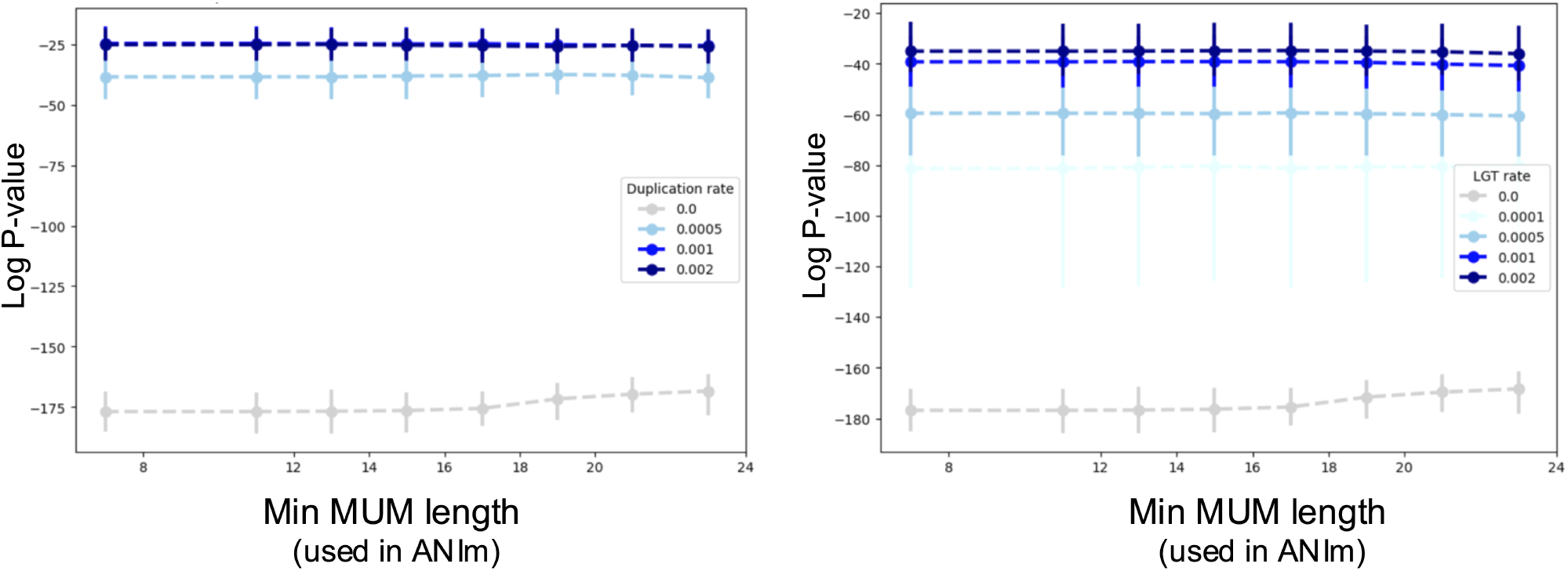
The correlation between ANIm∗alignment fraction and tree distance for different minimum MUM length. The alignment fraction here refers to the amount after removing duplicates using delta-filter -1 to keep only 1-1 alignments. In the last row we reported the alignment fraction before filtering. The simulated results show that the improvement for distant species achieved by weighted ANIm with the alignment fraction does not hold when there is high amount of duplications or LGT.

**Supplementary Figure 7:**
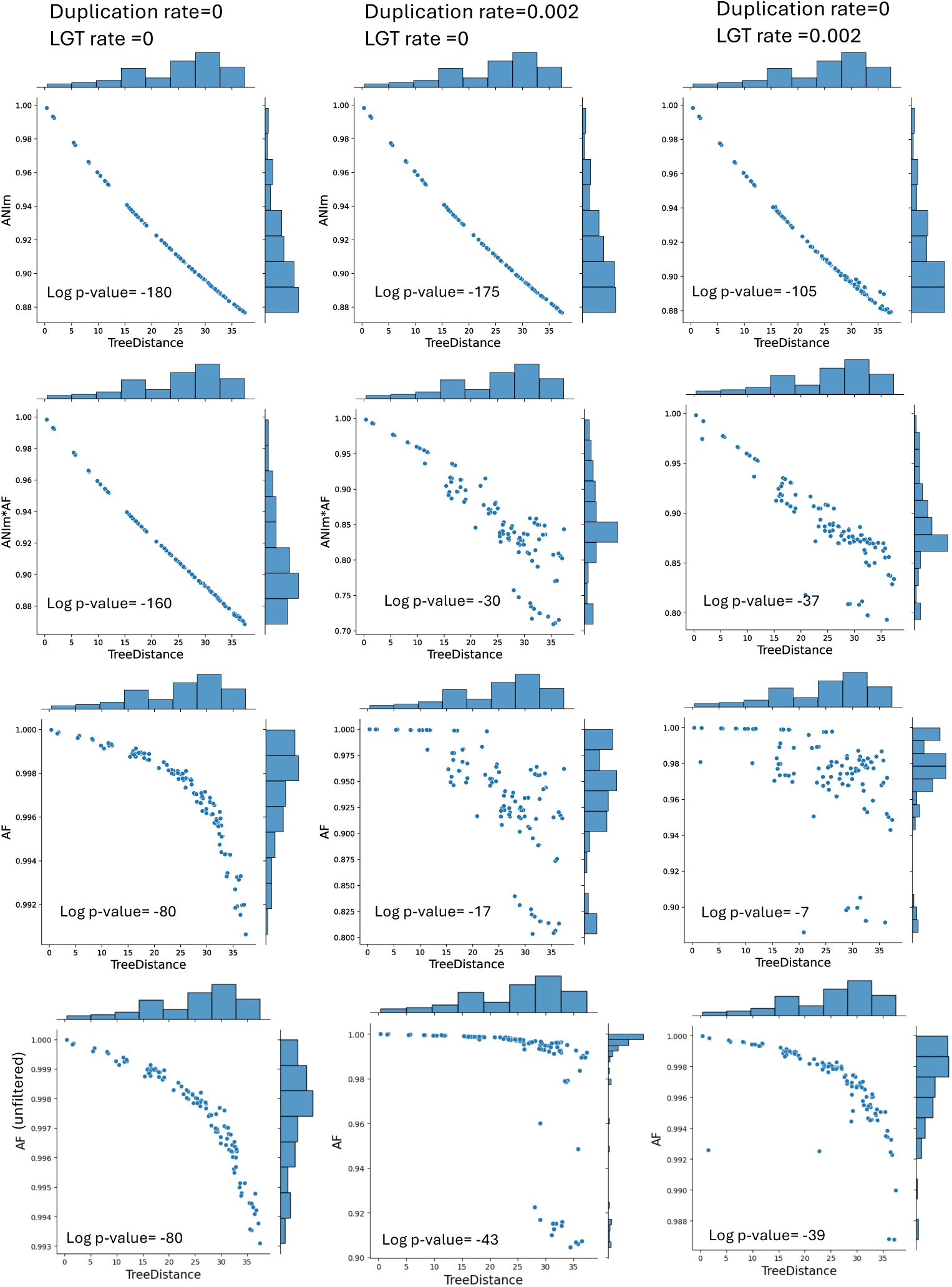
The correlation between ANIm∗alignment fraction and tree distance is impacted by the fact that *AF* is not well correlated with distance when there is higher LGT or duplication. The reported log p-values in the figure are based on the Spearman correlation test. Each point is one of the 105 pairs for 15 simulated genomes. The last two rows correspond to alignment fraction after and before filtering using delta-filter to find 1-to-1 alignments. In all figures, NUCmer was executed in MUM mode.

**Supplementary Figure 8:**
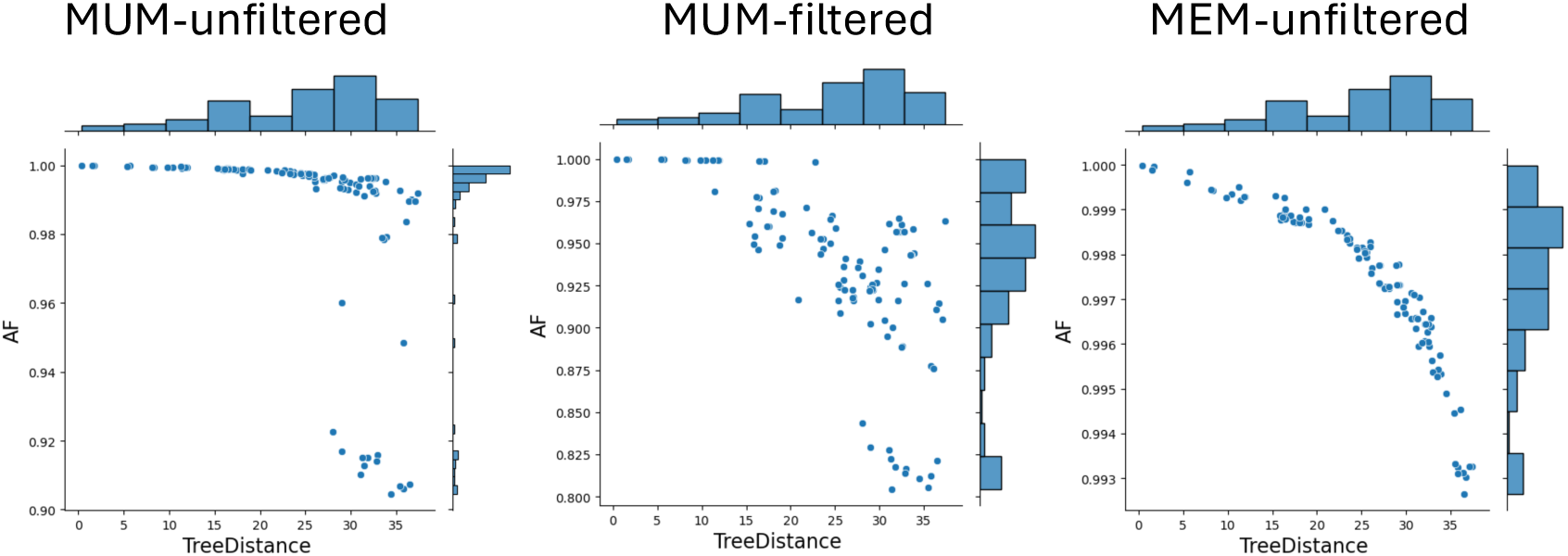
The impact of using MEM (maxmatch) instead of MUM with NUCmer in ANIm. Keeping only 1-to-1 alignments (MUM-filtererd) using delta-filter also decreases the alignment fraction.

**Supplementary Figure 9:**
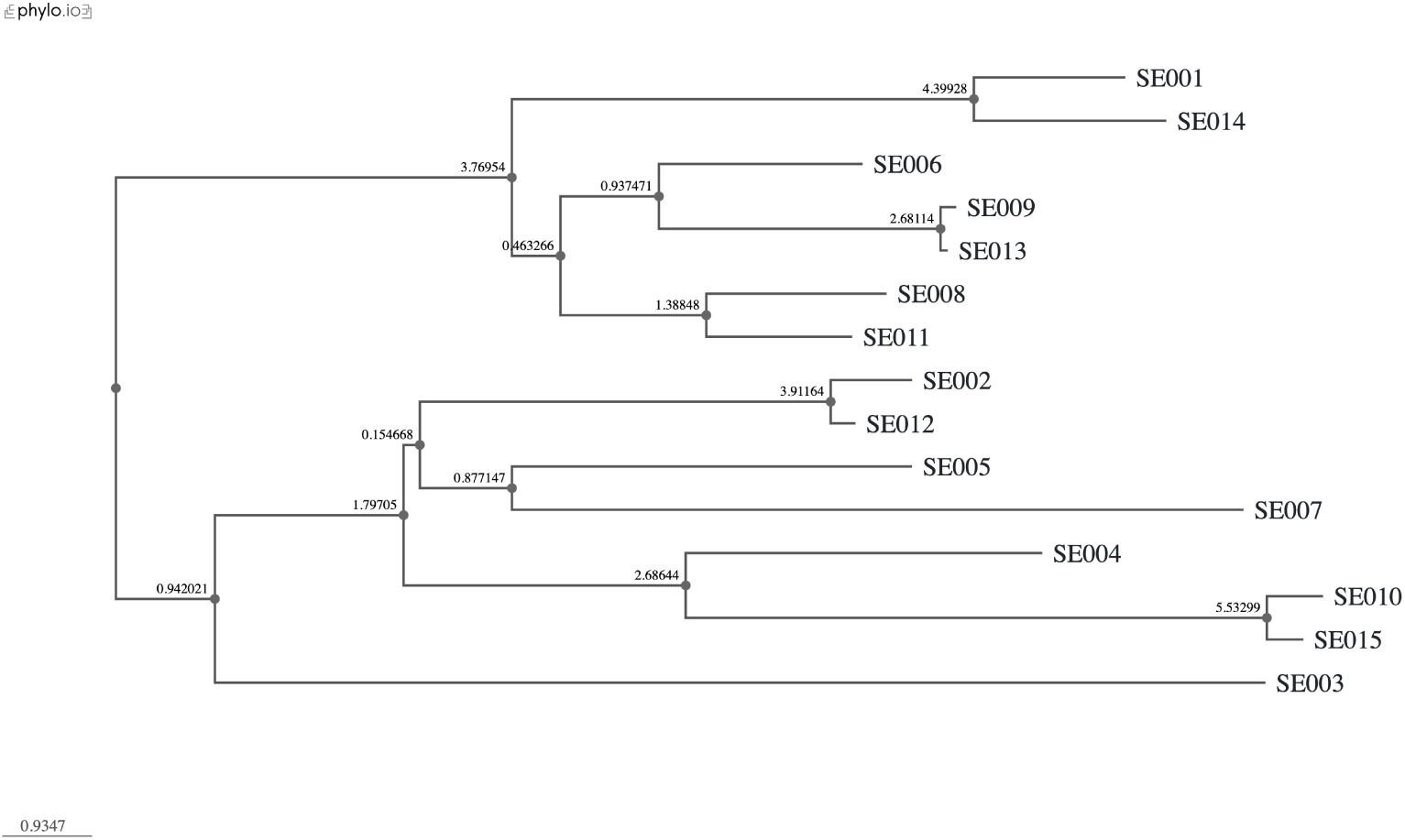
An example tree output of the ALF simulator with Mutation Rate of 10. The tree includes 15 species (SE001..SE015).

**Supplementary Figure 10:**
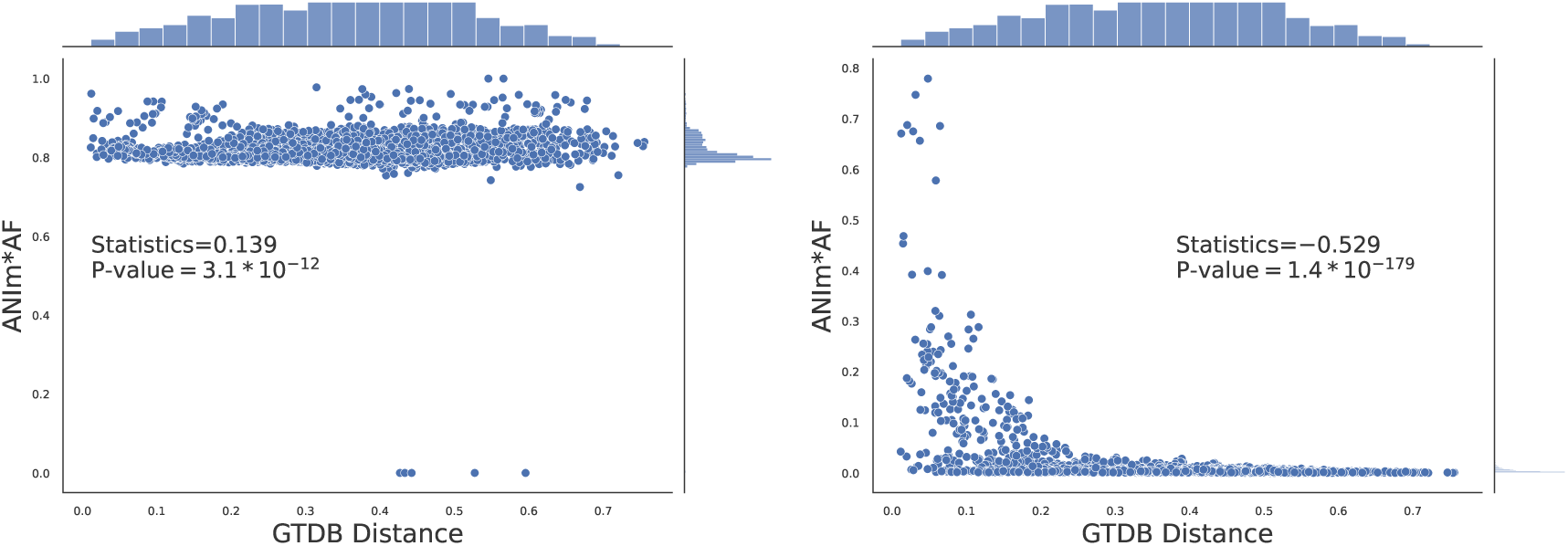
Impact of weighting ANIm with alignment fraction (AF) for distance calculation in Cyanobacteraia, similar to Figure 8 but here GTDB tree is used instead of NCBI tree. Here, 71 species are considered and 13 species were discarded due to incompatibility of species names between GTDB and NCBI taxonomy. The same pattern is observed with more pronounced p-values.

**Figure 10:**
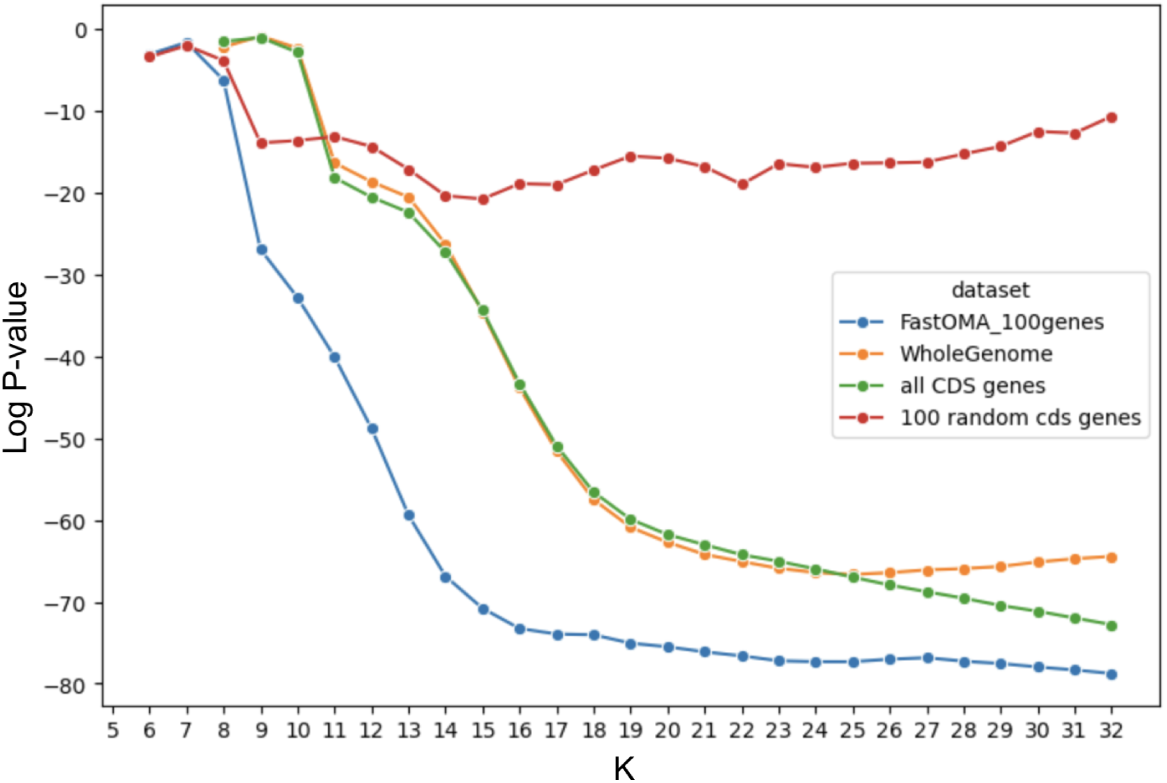
The log p-value of Spearman correlation test between the Jaccard index and distance on the genome taxonomy database (GTDB) tree (the lower, the better). This shows the impact of using different genomic regions in distance calculation for the clade 0 Bacillales A2. We considered k-mers found from the whole genome, all coding sequences (CDS), 100 random CDS genes, or 100 orthologous genes. Orthologous genes showed a stronger rank correlation with the GTDB tree distance compared to the whole genome using k-mers.

**Supplementary Figure 11:**
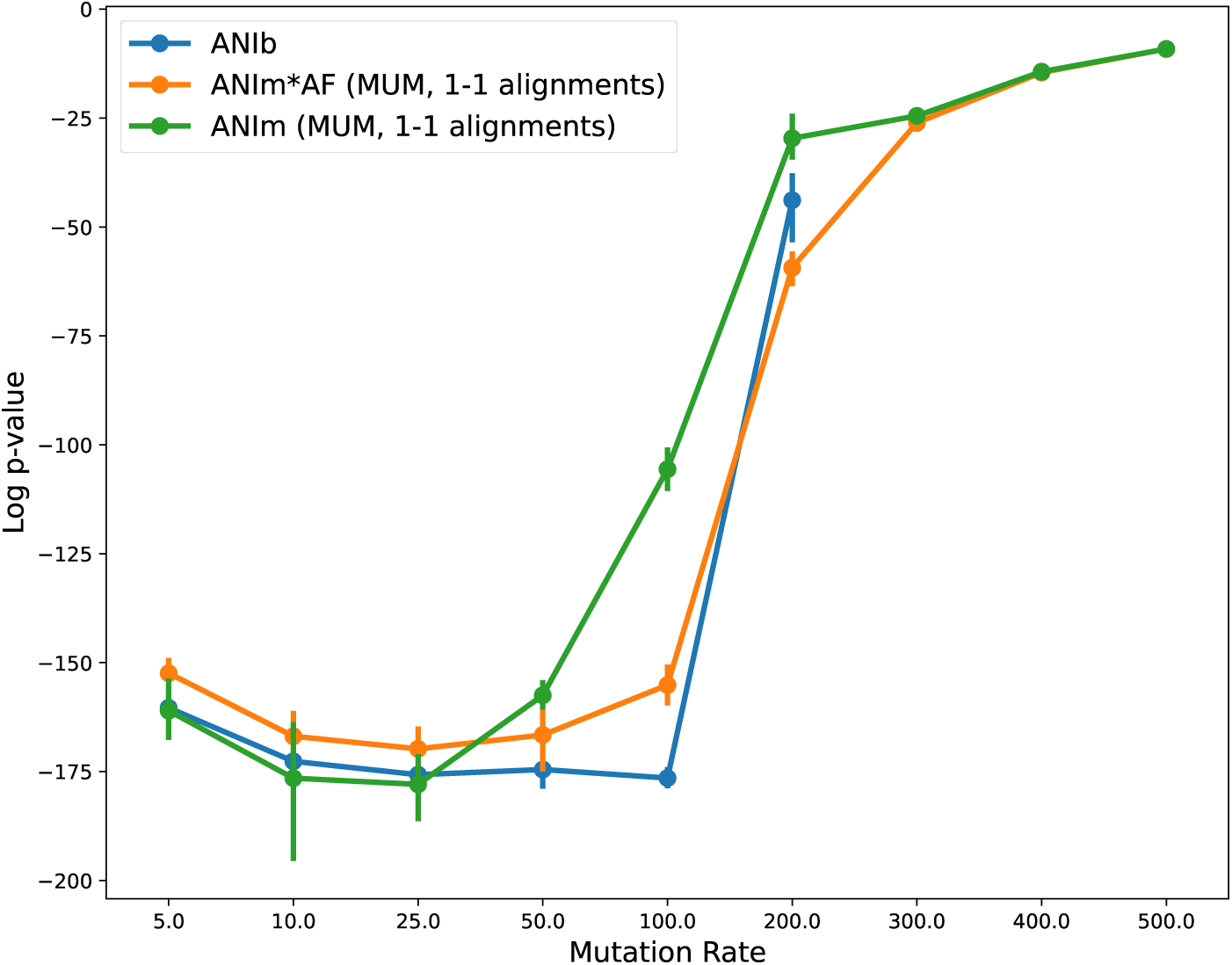
Comparing ANIb with ANIm and *ANIm* ∗ *AF* . Note that when mutation rate is very high (≥ 200), there is not much homology that can be detected by BLAST. BLAST output is empty for these cases when the e-value threshold was *_e_−*15.

**Supplementary Figure 12:**
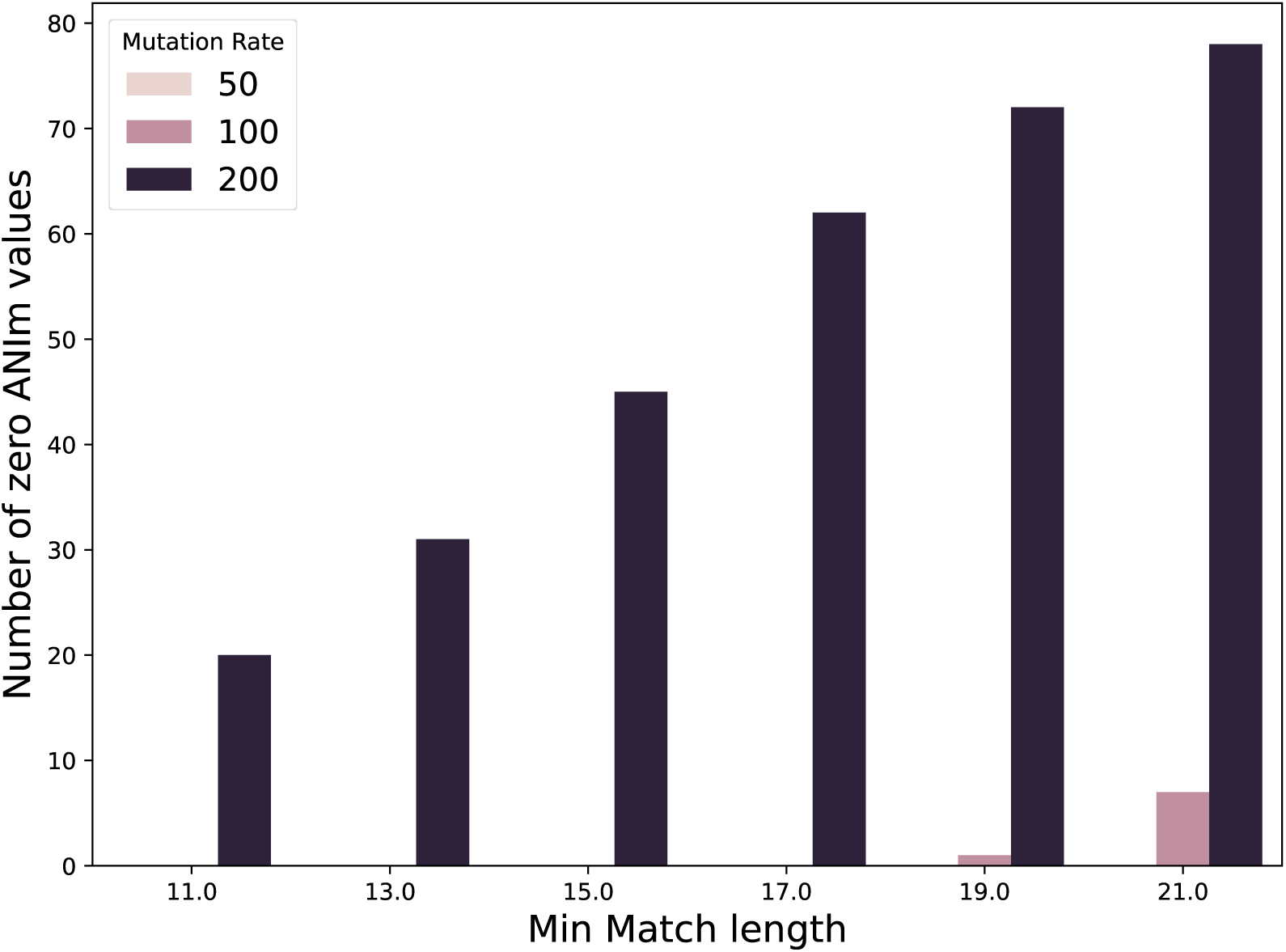
Number of zero ANIm values increases by mutation rate (more divergent dataset), affecting the power of ANIm. This could be mitigated by decreasing the minimum match length, ultimately improving rank correlation between ANIm and distance tree (e.g. in **Supplementary** Figure 3).

